# Functional avidity of anti-B7H3 CAR-T constructs predicts antigen density thresholds for triggering effector function

**DOI:** 10.1101/2024.02.19.580939

**Authors:** Marta Barisa, Elisa Zappa, Henrike Muller, Rivani Shah, Juliane Buhl, Benjamin Draper, Courtney Himsworth, Chantelle Bowers, Sophie Munnings-Tomes, Marilena Nicolaidou, Sonia Morlando, Katie Birley, Clara Leboreiro-Babe, Alice Vitali, Laura Privitera, Kyle O’Sullivan, Ailsa Greppi, Magdalena Buschhaus, Mario Barrera Román, Sam de Blank, Femke van den Ham, Brenna R. van ‘t Veld, Gabrielle Ferry, Laura K. Donovan, Louis Chesler, Jan Molenaar, Jarno Drost, Anne Rios, Kerry Chester, Judith Wienke, John Anderson

## Abstract

Chimeric Antigen receptor T cell (CAR-T) treatments for solid cancers have been compromised by limited expansion and survival in the tumor microenvironment following interaction with antigen-expressing target cells. Using B7H3 as a model antigen with broad clinical applicability, we evaluated the relationship between the antibody/antigen affinity of three clinical candidate binders and the three following functional characteristics: functional avidity, prolonged cytotoxicity in tumoroid re-stimulation assays, and *in vivo* anti-tumoral responses. BEHAV3D video-microscopy assessed distinct CAR-T cell behaviors at single cell resolution. T cell exhaustion did not dictate effector function. Rather, we demonstrated a threshold avidity of CAR-T / tumor cell interaction, characterized by longer cumulative CD8^+^ CAR-T / tumor target interaction times, and required for adequate CAR-T cell expansion to result in sustained tumor control upon re-challenge. These results provide new insights into design of CAR-T cells for antigen-dim cell targeting, and avoidance of antigen-dim tumor relapse.

## Introduction

Genetic modification of a cancer patients’ T cells with Chimeric antigen receptors (CAR-T cells) for adoptive transfer immunotherapy is a rapidly expanding clinical and scientific field. Clinical studies targeting hematological malignancy using adoptively-transferred CAR-T cell therapy have resulted in FDA approvals of CAR-T products for the treatment of leukaemia, lymphoma and myeloma ^1^. In contrast, far fewer durable clinical responses attributable to CAR-T cells are reported in solid cancers despite considerable efforts^2^. A major reason for the relative failure of CAR-T cell adoptive transfer in the solid cancer setting is suboptimal T cell functionality, evidenced by a lack of T cell proliferation and persistence. Reasons for this failure can likely be distilled down to inhibitory effects of the tumor microenvironment and inherent failure of CARs to provide optimal signals to the T cell upon target engagement.

CARs comprise an antigen-sensing ectodomain, which is typically a single chain variable fragment (scFv) of an antibody, and endodomains that are an amalgamation of ITAM-containing T cell receptor (TCR) signalling domains, most typically the ζ-chain of CD3, and co-stimulatory domains. CAR-T cells thereby commandeer the MHC-unrestricted antigen specificity of a monoclonal antibody and combine it with T cell signalling in a single molecule. The triggering of T cell activity is dependent on the formation of an effective immune synapse with an antigen-positive target cell. Once a productive synapse is formed, signalling occurs through engagement of the CD3ζ and co-stimulatory moieties with proximal signalling molecules, leading to T cell effector functions and a pro-inflammatory response.

For example, structural features of the CAR that influence persistence of effector function include hinge and transmembrane components, and choice and orientation of co-stimulatory endodomains. Moreover, certain scFv binders which signal in the absence of antigen contribute to excessive signalling and subsequent T cell dysfunction^3,4^. Studies in CD19-expressing leukaemia have shown that scFv affinity can affect the strength and duration of the immune synapse with consequences for subsequent disease control^5^. The quality and quantity of immune synapse formation are influenced by binder affinity (both on– and off-rate), target antigen density and CAR expression density. Functional avidity (strength of association between a CAR-T cell and its cellular target), is similarly governed by both affinity of interaction and expression levels of CAR and antigen in the interacting cells. A detailed understanding is lacking of how best to select an optimal CAR scFv for solid tumor-targeting CAR-T cells, and how scFv affinity and avidity impact on sustained functionality in solid tumors.

To address these challenges, we chose a broadly expressed solid tumor cancer antigen that is targeted in a number of current CAR-T clinical studies in solid tumors. B7H3 is expressed on solid and hematological cancer in the adult and pediatric settings^6,7^. A member of the immunoglobulin superfamily and closely related to PD-L1, B7H3 is enriched on high-grade tumors and expressed at low levels on healthy tissue^8–13^. There are several B7H3 binders that have been or are being progressed to evaluation in clinical trials including MGA271, 376.96 and TE9 ^7,9,14^. To specifically evaluate the impact of the scFv binding properties on CAR-T function we cloned these three different binders into identical viral backbones. All constructs had a 2^nd^ generation 28ζ endodomain and CD8 hinge and transmembrane format, and hence we set out to determine generalizable relationships between binder properties with CAR-T function rather than evaluate the binders in the context of their optimized formats for clinical studies.

Contrary to our expectations based on preceding literature^5,15–17^, we found that CAR-T incorporating either of two scFv’s that formed high avidity interaction with B7H3-expressing cells mediated superior anti-tumor functionality compared to a low avidity scFv. Both high and low avidity interactions enabled acute cytotoxicity and cytokine production, but only high avidity interactions led to sustained CAR-T proliferation and consequent activity in the re-challenge setting. These differences in scFv-driven functional potential were most evident against B7H3^med^ or B7H3^dim^ targets, the consequence of which was emergence of antigen-dim escape variants with the low avidity binder. We postulate that, in order to mediate tumour control, B7H3 CAR-T cells need to interact with their target cell with a supra-threshold avidity, sufficient to drive initial activation and proliferation.

## Results

### MGA271, 376.96 and TE9 scFv’s mediate divergent CAR-T responses against in vivo models of solid tumours

MGA271^18^, 376.98^19^ and TE9^20^ scFv’s were cloned into a CD8-28ζ 2^nd^ generation CAR format containing a CD8_a_ hinge-transmembrane, and expressed in SFG γ-retroviral vector with an EF1_a_ promoter as described previously^20^. A compact epitope tag, RQR8^21^, was co-expressed from the construct to monitor CAR-T transduction efficiency (Figure 1A). Comparable transgene co-expression between the binders was validated by staining the CAR directly with B7H3-His protein (Supplementary Figure 1). Against LAN-1 and Kelly neuroblastoma targets, we observed enhanced CAR-T cytokine production for 28ζ compared to 4-1BBζ endodomain for all the binders, consistent with previous literature including our own data^7,20,22^, and which was concurrent with higher checkpoint receptor expression (Supplementary Figure 2). CD28 co-stimulation was, therefore, chosen over 4-1BB.

**Figure 1.**
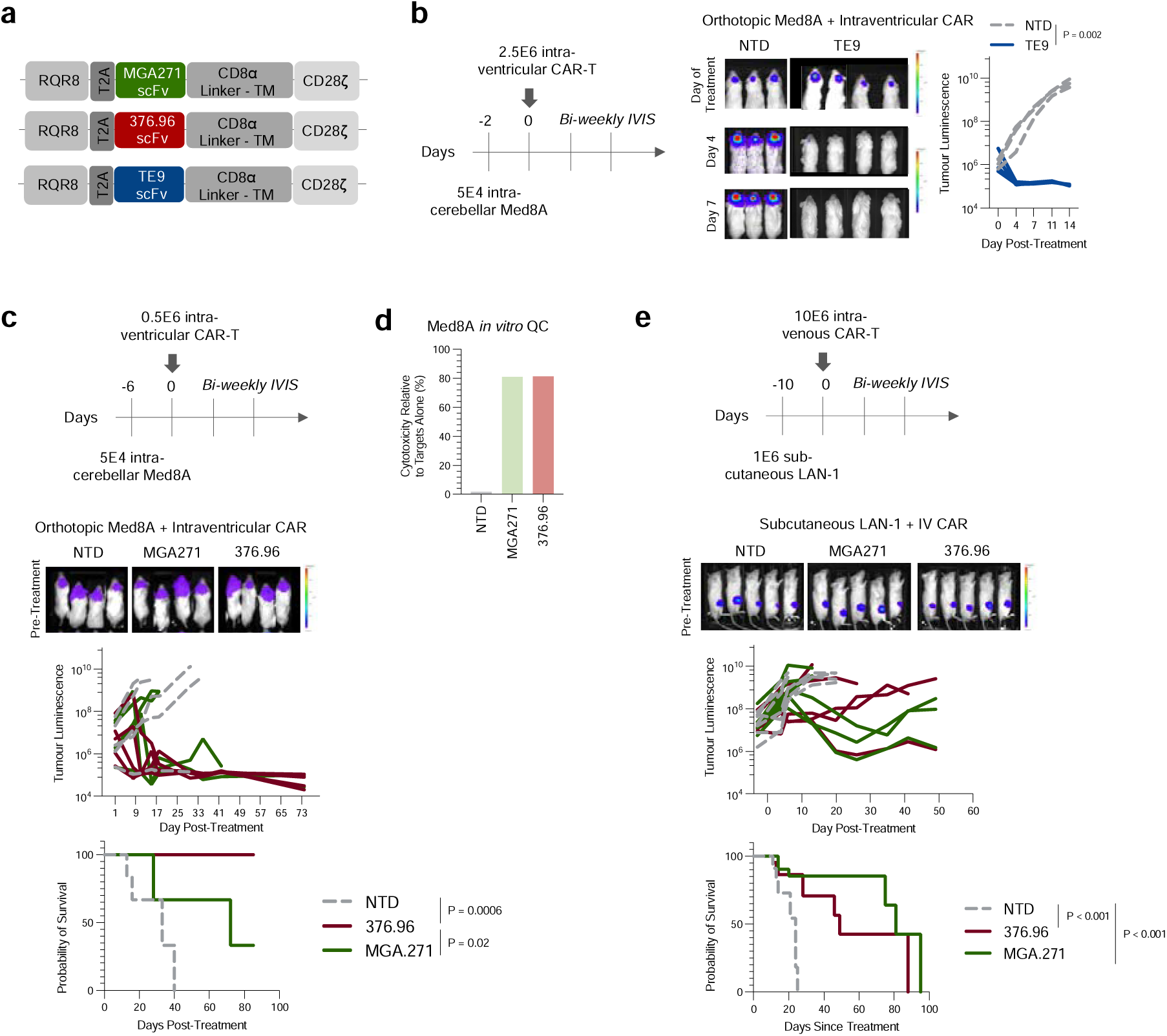
MGA271, 376.96 and TE9 scFv’s mediate divergent CAR-T responses against in vivo models of medulloblastoma and neuroblastoma. (**A**) Schematic of the retroviral constructs. (**B**) Orthotopic “non-stress” condition Med8A tumor model pilot in NSG mice; N=3 for NTD (non-transduced T cells), N=4 for TE9-CAR (tumor luminescence comparison between groups by Two-Way ANOVA). (**C**) Orthotopic “stress condition” Med8A tumor model in NSG mice; N=4 mice for all conditions (Kaplan-Meyer comparison by Log-rank Mantel-Cox test). (**D**) Overnight CAR-T cytotoxicity assay run in-parallel to the *in vivo* study from panel (**C**) (E:T ratio 1:10, N=1). (**E**) Subcutaneous LAN-1 tumor model in NSG mice. N=5 mice for all conditions (Kaplan-Meyer curve comparison by Log-rank Mantel-Cox test).

We first evaluated TE9 *in vivo* performance in an orthotopic, B7H3-positive^23^ Med8A model of medulloblastoma in which NOD.Cg-Prkdc scid Il2rg tm1Wjl /SzJ (NSG) mice received intraventricular CAR-T cells 48 hours after orthotopic cerebellar tumor engraftment. Used in these non-stress conditions, TE9-28ζ fully cleared the tumors within a week of CAR-T infusion (Figure 1B). Next, we evaluated the binders MGA271 and 376.96 in the same model but in increased stress conditions of longer tumor engraftment and lower CAR-T cell dose. Here the 376.96-28ζ treatment was curative (Figure 1C) whilst MGA271-28ζ did not significantly improve survival compared with non-transduced (NTD) T cells. This was unexpected, as both CAR-T products were equivalently cytotoxic against Med8A in a single challenge effector: target (E:T) ratio of 1:10 co-culture (Figure 1D). In a repeat experiment of the stress condition Med8A model, TE9-28ζ CAR-T efficacy matched that of 376.96-28ζ CAR-T, with both binders effecting more cures than MGA271-28ζ CAR-T (Supplementary Figure 3).

We have previously shown that MGA271, 376.96 and TE9 28ζ scFv CAR-T are equivalent in short-term single challenge cytotoxicity assays against LAN-1 neuroblastoma cells, and that TE9-28ζ CAR-T significantly enhance survival in a difficult-to-treat subcutaneous LAN1 NSG model^20^. We, therefore, evaluated MGA271 and 376.96-28ζ CAR-T in the LAN-1 *in vivo* model (Figure 1E). Intriguingly, in contrast with the Med8A model where differences between binders were apparent, 376.96 and MGA271 binders performed similarly against LAN-1 tumors, extending survival similarly to what was to described for TE9-28ζ^20^. Taken together, different *in vivo* models establish that B7H3 CAR-T functionality can be affected by scFv selection, but in a manner that is dependent on the choice of tumor target.

### MGA271 CAR-T have a lower avidity than 376.96 and TE9 CAR-T for B7H3-expressing tumor cells

Despite achieving similar high transduction efficiencies with all three binder CARs (Supplementary Figure 1), we noted that MGA271 CAR-T staining with B7H3-his protein was dimmer than that of matched 376.96 and TE9 CAR-T (Figure 2A, Supplementary Figure 4A). We hypothesized that the strength of the different scFvs’ binding to target, and its impact on strength of immune synapse might correlate with the differences observed between binders in the Med8A and LAN-1 *in vivo* studies. To test this, we evaluated the strength of cell-to-cell interaction for each of these CAR-T cells with Med8A, LAN-1, SupT1 wild type (SupT1-WT), and SupT1 transduced with 4-Ig human B7H3 (SupT1-B7H3^hi^), representing a range of antigen expression (Figure 2B). To avoid differences in transduction efficiency biasing avidity measurement, CAR-T of all scFv’s were diluted with donor-matched NTD T cells to standardize expression of the RQR8 marker gene (Supplementary Figure 4B).

**Figure 2.**
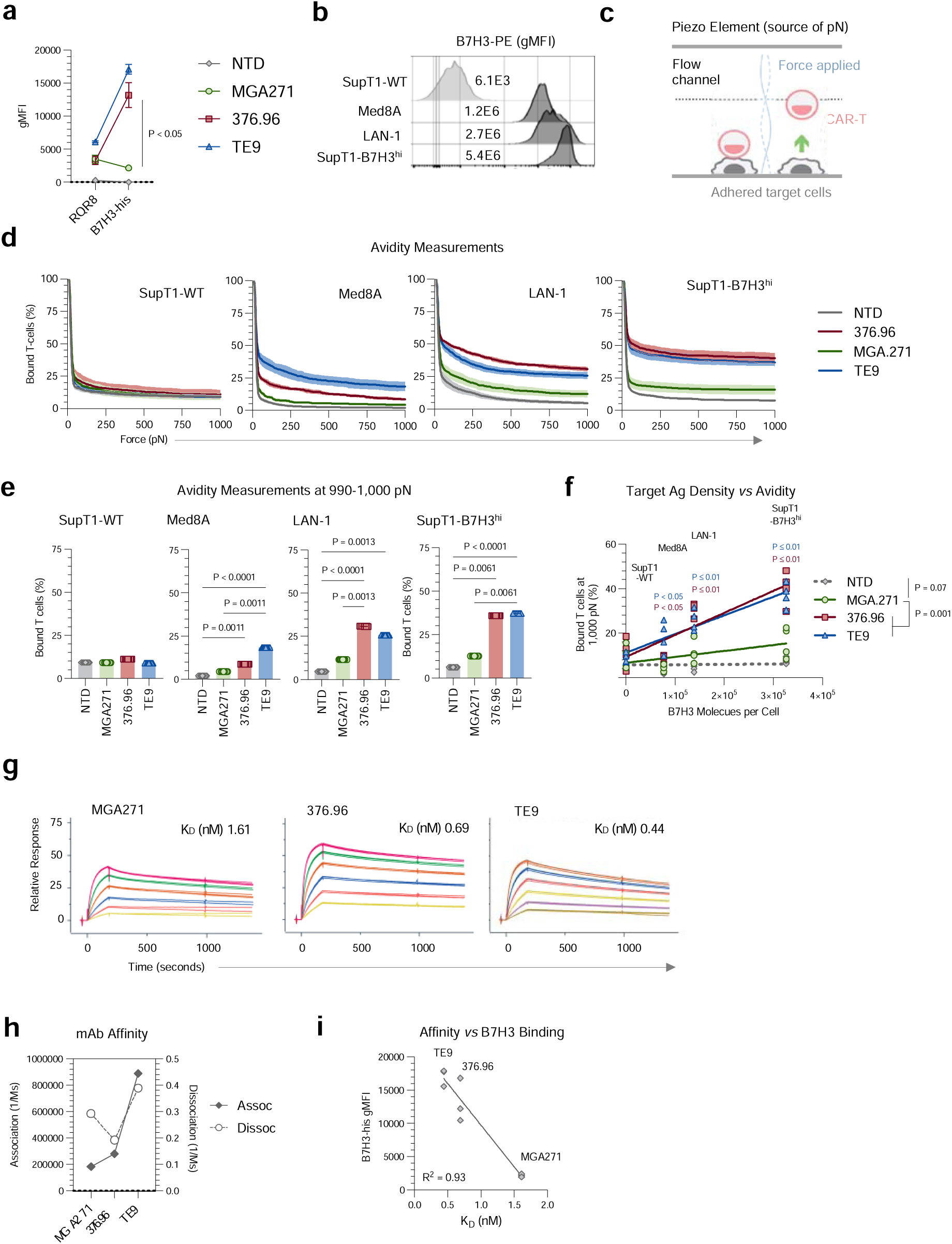
376.96 and TE9 but not MGA271 CAR-T form high avidity interactions with tumor cells in a B7H3 density dependent manner. (**A**) Cryopreserved, thawed CAR-T cell surface transgene expression was detected using flow cytometry with clone QBEND10 mAb to detect RQR8 marker gene and histidine-x6 tagged B7H3 protein (B7H3-his) + anti-his mAb to detect the CAR (mean ± SEM, N=3 donors, Two-Way ANOVA). (**B**) Target cells were stained with anti-B7H3-PE mAb and analyzed by FACS. Shown on the staggered histogram are respective geometric mean fluorescence intensities (gMFI) of B7H3-PE. (**C**) Principle of Lumicks z-Movi avidity measurement platform. (**D**) CAR-T detachment plotted against force in pN applied to the T cell / target interaction. The line indicates the mean T cell attachment of 4 independent donors across 3 experimental replicates whilst shaded area represents SEM. (**E**) Avidity as represented by attachment at 990-1,000 pN applied force: N=4 donors each with 3 experimental replicates, One-Way ANOVA. (**F**) Bound CAR-T (%) at 1,000 pN plotted against QuantiBright defined B7H3 molecules per cell. The lines are linear regressions of the means of 4 individual donors (Two-Way ANOVA). (**G**) Binder affinity for B7H3 (4-Ig) was measured using whole antibodies in IgG1 format using Biacore Surface Plasmon Resonance (SPR). (**H**) The association (‘on-rate’) and dissociation (‘off-rate’) rate constants for the different binders were plotted. (**I**) Binder affinity was plotted against the gMFI of B7H3-his protein staining of three independent donor CAR-T cell products.

Avidity measurements were carried out using a Lumicks z-Movi Cell Avidity Analyzer, which measures the force (in piconewtons, pN) required to detach a population of effector cells from their targets (Figure 2C). None of the CAR-T bound to B7H3^neg^ SupT1-WT cells, whilst all three bound B7H3^pos^ cells, albeit with different avidities (Figure 2D,E). TE9 and 376.96 scFv CAR-T consistently bound antigen-positive target cells with higher avidity than MGA271 or NTD cells.

To investigate the relationship between antibody-to-antigen binding properties, avidity and antigen target density, we measured the antibodies’ affinities, as well as target B7H3 molecules per cell in the respective target cell lines. Target cells with high B7H3 per cell overall formed higher avidity synapses with CAR-T, although the positive correlation between binding strength and antigen density was more pronounced for TE9 and 376.96 than MGA271 (Figure 2F). Binder affinity of binders in IgG1 format for 4-Ig B7H3 was measured using Surface Plasmon Resonance. TE9 had the highest affinity for B7H3 (0.44nM), similar to 376.96 (0.69mM) and higher than MGA271 (1.61nM) (Figure 2G, Supplementary Figure 5). It was of interest that TE9 had both the highest on-rate and the highest off-rate, whilst 376.9 had relatively slow on– and off-rate (Figure 2H). Since affinity is the ratio of off– and on-rates, dissociation constant (K_D_) values of TE9 and 376.9 were similar. Binder affinity for B7H3 correlated with the intensity of staining of CAR-T cells by recombinant B7H3 protein (Figure 2I).

### Cytotoxic capacity of low-avidity MGA271 CAR-T deteriorates upon rechallenges with B7H3^dim^ but not B7H3^bright^ tumors

To examine the impact of CAR-T binding avidity on effector function, the different constructs were assessed against solid tumor target cells that express a range of B7H3 densities. We selected five different B7H3^pos^ target types which were either cell lines or primary neurosphere cultures (defined hereafter as ‘tumoroids’), derived from pediatric solid tumors with a broad range of B7H3 antigen densities as determined using QuantiBright beads (Figure 3A).

**Figure 3.**
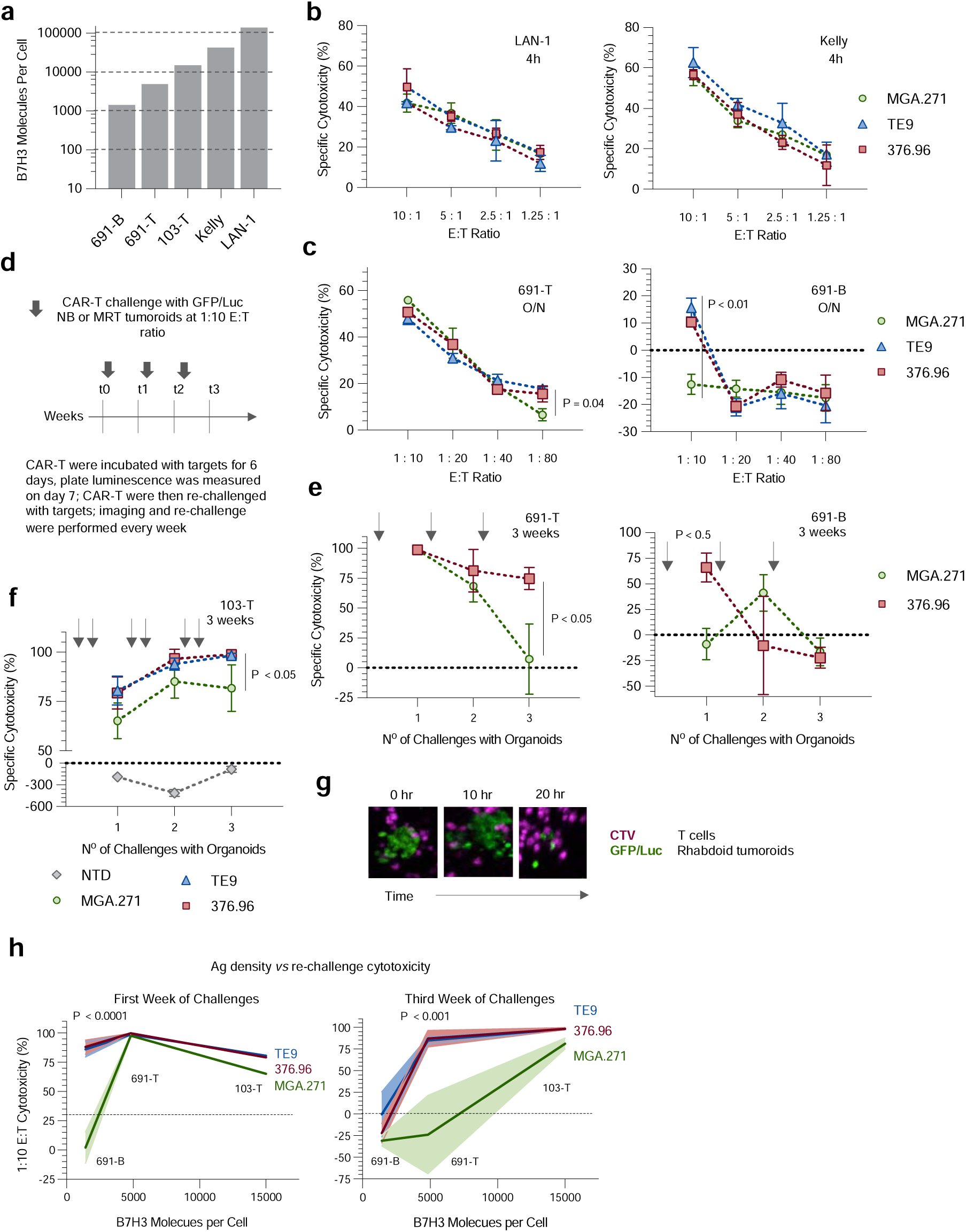
TE9 and 376.96, but not MGA271 CAR-T, respond to B7H3 ultra-dim tumor targets in serial re-challenge cytotoxicity assays. (**A**) Number of B7H3 molecules/cell (QuantiBright flow cytometry assay) for range of targets. (**B**) CAR-T cell cytotoxicity against LAN-1 and Kelly neuroblastoma cell lines (4h ^51^Cr release assay). ‘Specific cytotoxicity’ denotes cytotoxicity relative to non-transduced (NTD) T cell controls (mean ± SEM, N=3 donors, Two-Way ANOVA). (**C**) CAR-T cell cytotoxicity against GFP/Luc-engineered tumoroids by overnight luminescence-based assay. ‘Specific cytotoxicity’ denotes cytotoxicity relative to NTD T cell controls (mean ± SEM, N=2 independent donors across 3 technical replicates, Two-Way ANOVA). (**D-F**) Re-challenge assay timecourse (**D**), against neuroblastoma (**E**) and malignant rhabdoid tumor (MRT) organoids (**F**); gray arrows indicate re-challenge with tumor. In (**E**), data is mean ± SEM, N=2 independent T cell donors across 3 experimental replicates and ‘specific cytotoxicity’ denotes cytotoxicity relative to NTD T cell controls; in (**F**) ‘specific cytotoxicity’ denotes cytotoxicity relative to tumor alone controls plated in parallel, data is mean ± SEM, N=1 independent T cell donor (same as one of the donors used to generate data for panel **E**) across 3 experimental replicates. (**G**) Representative microscopy images of 103-T MRT tumoroid being killed by CAR-T over time. (**H**) 1:10 E:T ratio cytotoxicity for the first and third week challenge from the assays showed in panels (**E** and **F**) plotted against antigen density (Quantibright); line = mean cytotoxicity, shaded area = SEM.

CAR-T cytotoxicity was first evaluated at a range of E:T ratios in single-challenge short-term assays. All binders displayed equivalent effective cytotoxicity against B7H3^hi^ neuroblastoma cell lines LAN-1 and Kelly (Figure 3B), but relative failure of the lower affinity MGA271 binder was seen when targeting the lower B7H3 antigen density neuroblastoma tumoroids at low E:T ratios (≤1:10) (Figure 3C).

Capacity of CAR-T constructs to maintain killing on repeat encounter with target cells is essential for their clinical utility. Therefore, CAR-T were re-challenged over a four-week co-culture as illustrated in Figure 3D. Allowing the CAR-T 6 days to effect cytotoxicity, 691-T tumoroids were killed by both low-avidity CAR MGA271 and representative high-avidity CAR 376.96 following the first challenge (Figure 3E). However, MGA271 killing dropped drastically after re-challenge and by the third challenge no MGA271-mediated 691-T killing was observed, while high avidity 376.96 CAR-T maintained tumor control, similar to TE9 CAR-T in an independent experiment (Supplementary Figure 6). Using 691-B tumoroids which have lower antigen density, the high avidity binders 376.96 and TE9 completely eliminated targets after a single challenge whilst MGA271 showed no cytotoxicity. All CAR constructs failed against 691-B upon re-challenge (Figure 3E, Supplementary Figure 6). A similar trend was observed for B7H3^med^ 103-T malignant rhabdoid tumoroids. All CARs were cytotoxic against 103-T, but the inferiority of MGA271 was the most pronounced upon challenge during the third and final week (Figure 3F,G).

Potential explanations for CAR-T failure in repeat stimulation assays is T cell exhaustion or differentiation to terminal effector cells. While we identified upregulation of exhaustion/activation receptors (including PD-1, TIM-3 and LAG-3) on all scFv CAR-T cells following challenge with neuroblastoma targets (Supplementary Figure 2 for Kelly and LAN-1 cell lines; Supplementary Figure 7 for 691 tumoroids), these were not consistently different between the different binders, and, therefore, did not appear to account for functional differences between the CAR products. There were similarly no consistent differences between binders in memory markers CD45RA and CCR7, nor activation markers CD25 and CD69 (Supplementary Figure 7). Collating data from the tumoroid co-cultures at 1:10 E:T ratio, the differences in cytotoxicity between binders at low antigen density was pronounced at both initial challenge and re-challenges, and is represented graphically in Figure 3H. At the lowest antigen density (∼1,000 molecules per cell for 691-B tumoroids), only the high avidity binders, TE9 and 376.96, were capable of initiating killing at first challenge but failed to sustain cytotoxicity at re-challenge. At ∼5,000 molecules per cell or more, all the binders were effective at initial challenge but only the two high avidity binders sustained killing over multiple rounds of stimulation (Figure 3H).

The data prompted us to hypothesize that at high antigen density, binders perform similarly because CAR-T signalling for all of them exceeds a threshold for proliferation and cytokine secretion to initiate tumour control. At low antigen density, high avidity interaction is required to exceed this threshold. Therefore, we next evaluated CAR-T effector functions other than cytotoxicity, including IL-2 and IFN-γ production, and CAR-T cell proliferation.

### High avidity interaction with target cells drives CAR-T proliferation and thereby sustained effector function

To compare activation profiles, we first measured CAR-T cytokine secretion in response to antigen-bright LAN-1 and Kelly neuroblastoma cell lines. Unexpectedly, despite having seen high and equivalent killing of the targets by all three CARs in short-term killing assays (Figure 3B), the lower avidity MGA271 produced significantly lower levels of IFN-γ and IL-2 than TE9 and 376.96 CAR-T (Figure 4A,B). MGA271 CAR-T cell numbers declined throughout the assay – in a manner that was indistinguishable from NTD T cells, whilst TE9 and 376.96 CAR-T proliferated at each challenge with targets (Figure 4C).

**Figure 4.**
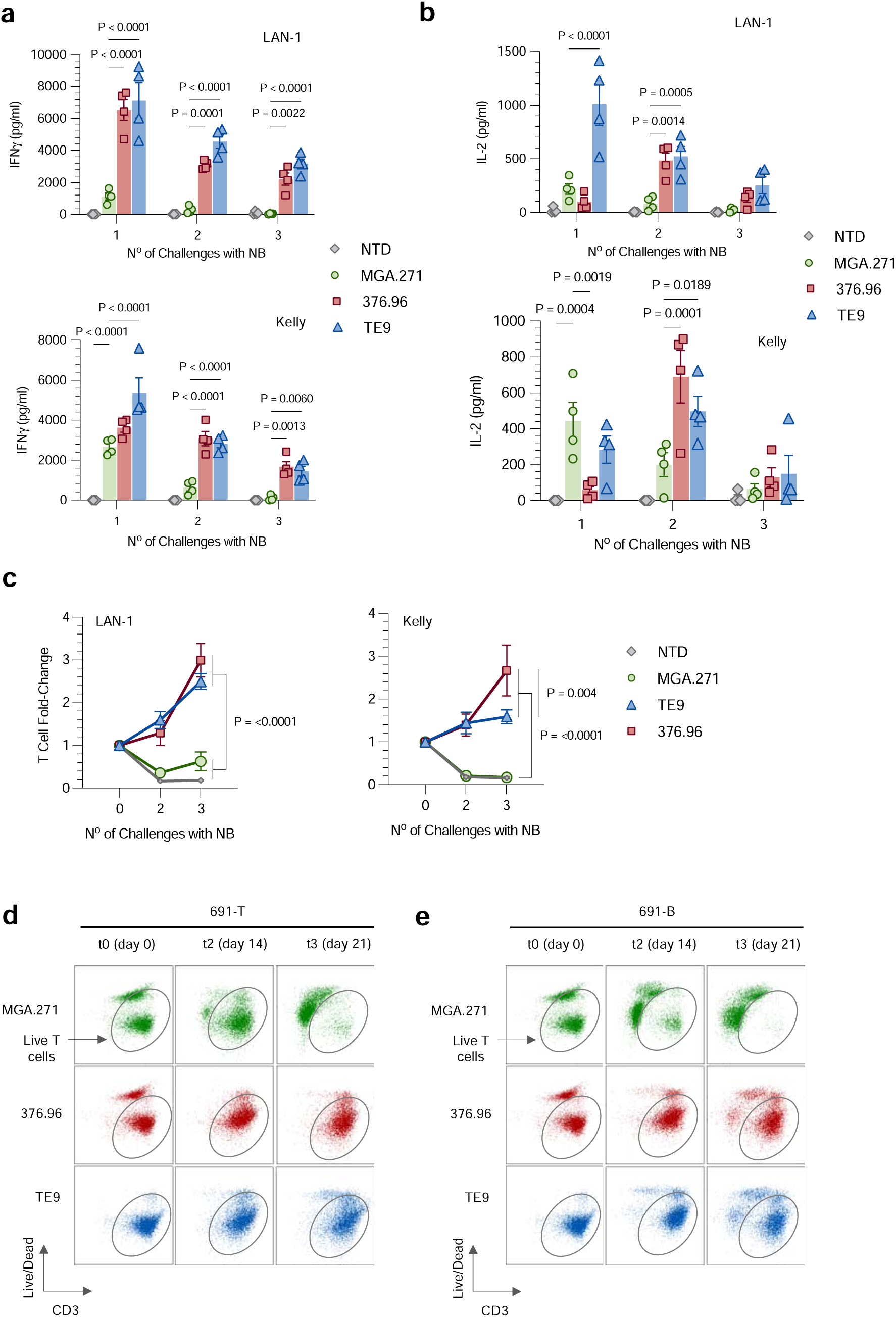
High avidity interaction with target cells drives CAR-T proliferation and resulting sustained effector function. (**A**,**B**) CAR-T cells were stimulated with irradiated LAN-1 or Kelly neuroblastoma cell lines at a 1:1 E:T ratio every 7 days; IFN-γ (**A**) and IL-2 (**B**) were measured in supernatants collected 24h after every stimulation (mean ± SEM, N=3 independent donors, Two-Way ANOVA). (**C**) T cell numbers in the same co-cultures were evaluated at each time point using Precision Count Beads and flow cytometry (mean ± SEM, N=3 independent donors, Two-Way ANOVA). (**D**) Relative T cell proportions measured using flow cytometry in 691-B and 691-T tumor co-cultures over three weeks of weekly tumor challenges at an E:T ratio of 1:10. Gated on singlet PBMC of harvested cultures, representative dot plots from one donor CAR-T are shown.

The pattern of CAR-T persistence and proliferation in the 691-T or –B neuroblastoma tumoroid re-challenge assays (Figure 4D and E) mirrored the differential cytokine responses of the different binders. MGA271, but not TE9 and 376.96, numbers declined steadily over time – even in conditions where the MGA271 CAR-T had initially mediated high level killing at first challenge such as was seen against 691-T in Figure 3C and E.

To evaluate whether B7H3 CAR-T responses to brain tumor targets are similar, we identified three different medulloblastoma models with a range of antigen densities (Figure 5A). All were B7H3^med^ or B7H3^med-hi^. CAR-T cytotoxicity upon single challenge with these targets was equivalently high across all scFv’s at a range of E:T ratios (Supplementary Figure 8). As was the case with the neuroblastoma targets, however, there were differences in antigen induced CAR-T proliferation between the binders. Low avidity MGA271 proliferated significantly less than TE9 or 376.96 CAR-T, at a level that was not significantly greater than that of NTD T cells (Figure 5B). Proliferation for all CARs was proportionate with antigen expression, and the required B7H3 threshold for driving an increase in T cell numbers was higher for MGA271 than TE9 and 376.96 (Figure 5C). Linking CAR-T proliferation to avidity measurements from Figure 2 revealed that for both Med8A and LAN-1 targets, avidity appeared predictive of subsequent T cell proliferation (Figure 5D).

**Figure 5.**
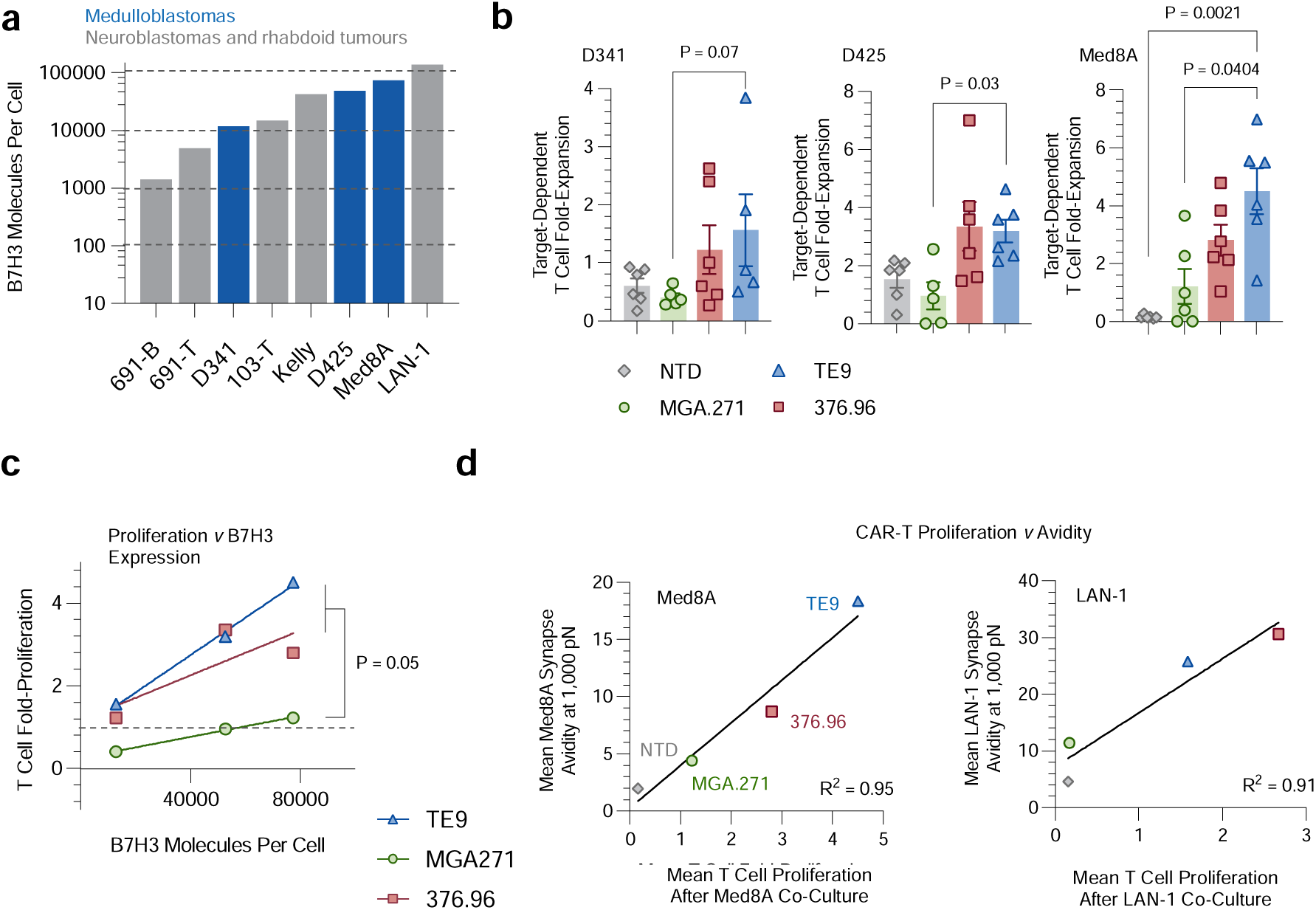
High avidity interactions drive CAR-T proliferation and cytotoxicity in brain tumor models. (**A**) B7H3 molecules per cell were quantified using a QuantiBright flow cytometry assay; brain tumor targets in blue, neuroblastoma and rhabdoid tumor targets in gray. (**B**) T cell proliferation in response to live brain tumor challenge at a 1:1 E:T ratio was quantified after 6 days of co-culture (mean ± SEM, 3 independent T cell donors across two experimental replicates, Kurskal-Wallis test). (**C**) T cell fold-expansion against the number of B7H3 molecules per target cell; dots represent mean proliferation of the different donors and replicates; lines are simple linear regression curves. (**D**) Mean fold-proliferation of CAR-T cells against Med8a and LAN-1 cells was plotted against the mean avidity at 1,000 pN for the two cell lines. The dots represent mean proliferation of the different donors and replicates; the lines are simple linear regression curves.

### High avidity, determined by ScFv affinity and B7H3 antigen density, drives CAR-T activation

To identify the threshold of B7H3 antigen density for binder-dependent CAR-T functionality within the context of the same target cell, we generated a cell line that expresses a range of B7H3 densities by transducing SupT1 cells with 4-Ig B7H3 (Figure 6A). The SupT1 cells were also transduced with a transgene encoding eGFP and luciferase to enable cytotoxicity evaluation by luminescence. Resulting SupT1-B7H3^range^ contained cells with range of B7H3 antigen densities from negative to ultra-high (Figure 6B). CAR-T were challenged with SupT1-B7H3^range^ cells at a 1:1 E:T ratio daily for a week to mimic continued exposure to tumor (Figure 6C).

**Figure 6.**
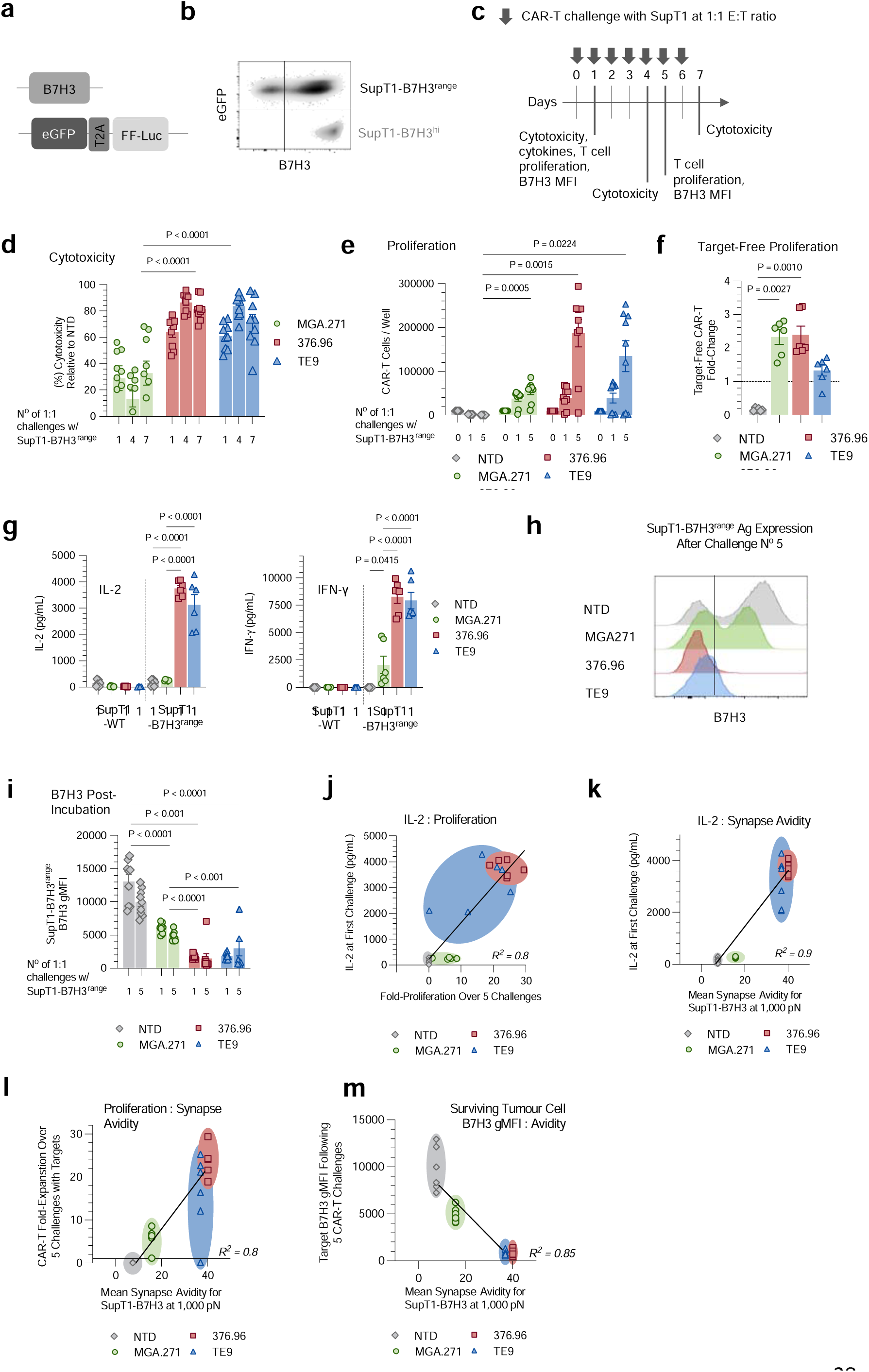
High avidity drives elimination of antigen-dim targets. (**A**) Constructs for generation of B7H3 and GFP/luciferase sublines of SupT1 cells. (**B**) Expression of B7H3 in the unsorted ‘SupT1-B7H3^range^’ and flow-sorted ‘SupT1-B7H3^hi^’ cells. (**C**) Rechallenge assay experimental schematic. For all subsequent data in the panels: N=4 independent donors across 4 experimental replicates. (**D**) Cytotoxicity was measured by luciferase signal as relative to tumor luminescence in the presence of donor-matched non-transduced (NTD) T cells (mean ± SEM, Two-Way ANOVA). (**E**) T cell numbers in the same assay following either a range of challenges with tumor targets or (**F**) target-free (mean ± SEM, Two-Way ANOVA). (**G**) Cytokine measurements in co-culture supernatants following a single challenge wit SupT1-B7H3^range^ targets at a 1:1 E:T ratio (mean ± SEM, Two-Way ANOVA). (**H**) B7H3 expression on targets was measured using flow cytometry following the fifth challenge. Shown are concatenated histograms that were generated by combining flow data adjusted for 10,000 SupT1 cells per sample. (**I**) The same SupT1-B7H3^range^ positivity for antigen as analysed in each sample separately (mean represents geometric mean fluorescence intensity (gMFI) ± SEM, Two-Way ANOVA). (**J**) Respective CAR fold-proliferation in response to 5 challenges with SupT1-B7H3^range^ targets plotted against IL-2 production following the initial challenge. (**K**) IL-2 production after first challenge plotted against the mean avidity for SupT1-B7H3^hi^ targets at 1,000 pN. (**L**) Proliferation after 5 challenges with targets was plotted against the mean synapse avidity that the respective CARs achieved when combined with SupT1-B7H3^hi^ targets at 1,000 pN. (**M**) SupT1-B7H3^range^ cell B7H3 gMFI after 5 challenges with targets was plotted against the mean synapse avidity that the respective CARs achieved when combined with SupT1-B7H3^hi^ targets at 1,000 pN. In (**J**-**M**) each dot represents an individual donor experimental replicate, the lines are simple linear regressions.

While all CAR-T cells exhibited some cytotoxicity against SupT1-B7H3^range^ cells, MGA271 killing of targets was consistently lower than that of TE9 and 376.96 CAR-T (Figure 6D). Of note, in contrast to previous targets assayed, MGA271 CAR-T kept up sustained, if low-level, killing even following 7 challenges with targets. None of the CAR-T killed control SupT1-WT cells (Supplementary Figure 9A). Substantial 376.96 and TE9 proliferation was seen as target challenge progressed, whilst more modest MGA271 proliferation was observed (Figure 6E). The antigen dependency of the proliferation was demonstrated by the lack of response in the presence of SupT1-WT targets (Supplementary Figure 9B) or unstimulated conditions (Figure 6F). TE9 and 376.96 CAR-T produced significantly more IL-2 and IFN-γ than MGA271 after the initial challenge with SupT1-B7H3^range^ targets (Figure 6G).

**Figure 7.**
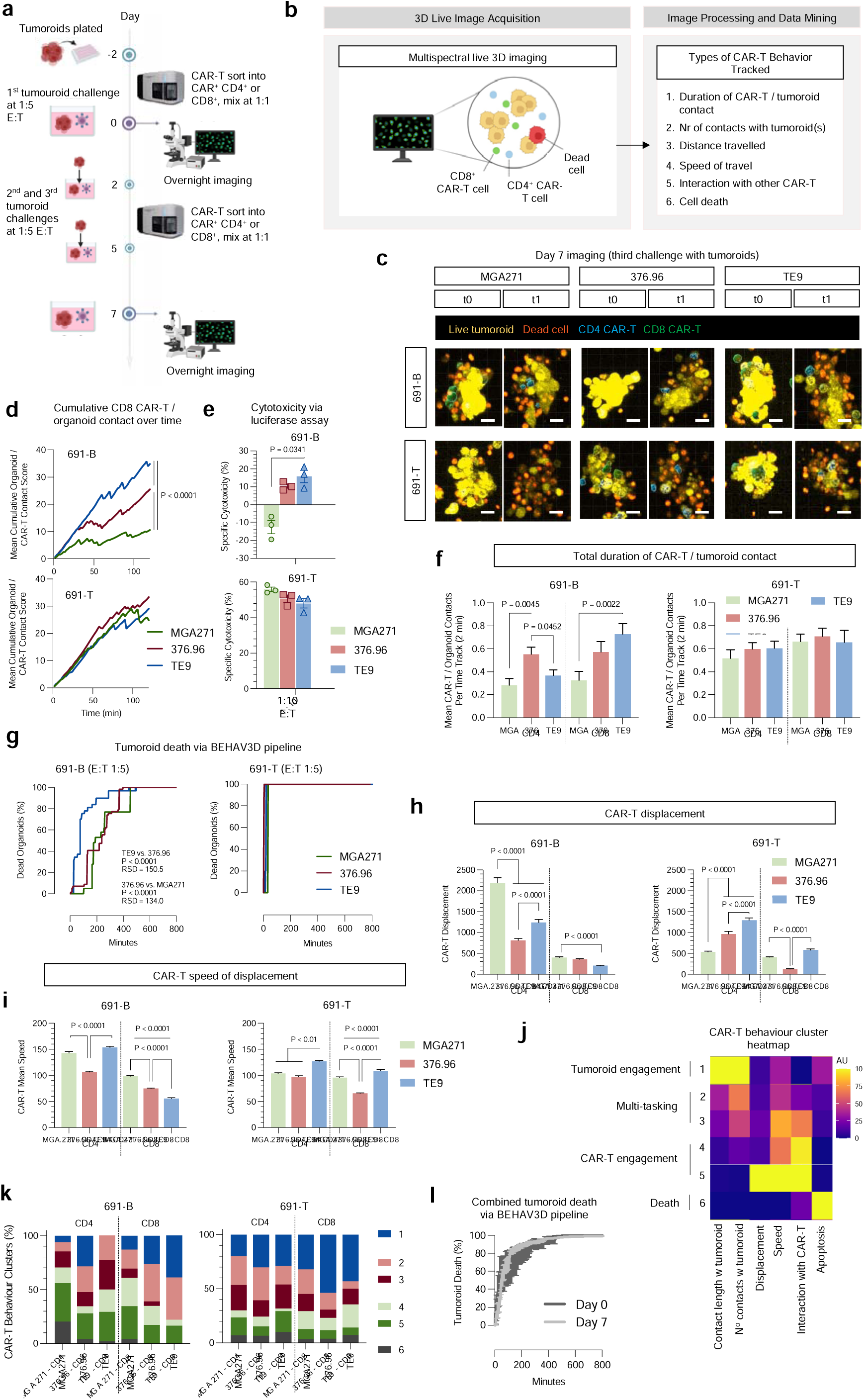
Higher CAR binder B7H3 avidity associated with increased CD8^+^ CAR-T / tumoroid contact duration and more rapid cytotoxicity against B7H3^ultra-dim^ tumoroids. (**A**,**B**) experimental overview. (**C**) Illustrative stills of tumoroid and T cell clusters at the start (day 0 = t0), and end (day 7 = t1) 800 minute imaging sessions. (**D**) CD8^+^ CAR-T cell cumulative contact time with 691-B and 691-T organoids tracked for the first 2h of the first tumor challenge on day 0. The line represents the mean cumulative tumoroid contact score of all the individual CAR-T cells that were successfully tracked for the full 120 min period (the number of cells tracked in each of the conditions was a mean 31.33±10.8; curves were compared using simple linear regressions). (**E**) CAR-T cells were incubated with tumoroids overnight at a 1:10 E:T ratio. ‘Specific cytotoxicity’ denotes tumor luminescence relative to matched non-transduced T cell (NTD) controls (mean ± SEM, N=3 independent experimental replicates, Kruskal-Wallis test). **(F**-**I)** Respective parameters were tracked for entire duration (800 minutes) of the first tumor challenge at day 0. (**F**) CD4^+^ and CD8^+^ CAR-T cell cumulative contact time with 691-B and 691-T organoids (mean ± SEM, N = 31.5 ± 10.6 cells per T cell condition, Two-Way ANOVA). (**G**) Cumulative tumoroid death (conversion from viability dye yellow to red) (a total of 398 incremental time-points were recorded to track tumoroid apoptosis over the assay; curve comparison was carried out using a Friedman test for P value and rank sum differences (RSD)). (**H**) CD4^+^ and CD8^+^ CAR-T displacement (mean ± SEM, N = 31.5 ± 10.6 cells per T cell condition, Two-Way ANOVA). (**I**) Mean speed of CD4^+^ and CD8^+^ CAR-T displacement (mean ± SEM, N = 31.5 ± 10.6 cells per T cell condition, Two-Way ANOVA). (**J**) Trends in CAR-T behavior reduced into 6 clusters using UMAP analysis of the T cell behavior across all the conditions observed in the initial tumor challenge at day 0 and represented as a heatmap. (**K**) The CAR-T behaviors broken down by T cell, binder and target type. (**L**) Pooled data from all CAR constructs against both targets comparing day 0 and day 7 challenge rate of organoid death.

CAR-T / tumor co-cultures were harvested after the 5^th^ challenge, and residual SupT1-B7H3^range^ expression of B7H3 was measured. Figure 6H shows concatenated B7H3 expression on GFP-positive SupT1 cells across the CAR groups. While TE9 and 376.96 CAR-T cultures contained few or no residual B7H3-positive targets, MGA271 CAR-exposed SupT1 cells maintained a range of B7H3 expression. Compared to the NTD T cell condition, MGA271 eliminated only the antigen-high SupT1 targets (Figure 6H,I).

We then aimed to determine the relationship between-T effector function (cytokine secretion and proliferation) and CAR avidity. CAR-T fold-proliferation over 5 challenges with targets correlated with IL-2 production at the time of the initial target challenge, though 376.96 CARs were more consistent initial IL-2 producers than TE9 CARs (Figure 6J). The avidity of CAR-T interaction with SupT1-B7H3^hi^ targets shown in Figure 2 correlated with both IL-2 production and proliferation in response to SupT1-B7H3^range^ cells (Figure 6K,L). A similarly strong but inverse correlation was observed between avidity of interaction and the antigen expression on residual SupT1-B7H3^range^ cells following serial re-challenge (Figure 6M). Taken together, the data indicate that for each cell target engaged by a standard CD28ζ CAR-T cell, there is a threshold of avidity required for sufficient cytokine and proliferation response to allow for expansion and effective cytotoxicity on target re-challenge. We next examined the impact of scFv on B7H3 CAR-T functionality at the single cell level.

### High ‘on-rate’ is associated with high CD8^+^ CAR-T / tumoroid contact duration and rapid cytotoxicity against B7H3^dim^ targets

BEHAV3D, a multispectral, 3D image-based platform, is designed to live-track the efficacy and mode of action of cellular immunotherapy at the single cell level ^24^. BEHAV3D was used to track anti-B7H3 CAR-T functionality over three challenges with 691-B and 691-T tumor targets. Cultures were imaged during the first (day 0) and third (day 7) challenge with targets, and the series of images then processed for CAR-T behavior using a bioinformatic pipeline, developed by Anne Rios and colleagues^24^ (the assay setup is illustrated in Figure 7A and B and Supplementary Figure 10). CAR-T cell behavior was tracked across six different parameters, shown in Figure 7B. Video compilations of day 0 and day 7 CAR-T / tumoroid co-cultures can be viewed in Supplementary Videos 1-24.

CAR-T/tumoroid interactions leading to tumoroid death was evident for all conditions (representative co-cultures images shown in Figure 7C). In analysis of behaviors against B7H3^dim^ 691-B tumoroids, striking and unexpected differences were seen between the binders, and between CD4^+^ and CD8^+^ T cells. Plotting the cumulative CD8^+^ CAR-T/tumoroid contact since initiation of the day 0 co-culture showed that TE9 and 376.96 CD8^+^ CAR-T spent significantly longer in contact with B7H3^dim^ 691-B tumoroids than MGA271 CD8^+^ CAR-T did (Figure 7D). A summation of total CD8^+^ CAR-T / 691-B contact time during the entire day-0 analysis confirmed this difference between binders (Figure 7F). This cumulative contact score correlated with the cytotoxicity of respective binder-CAR-T constructs against these B7H3^dim^ 691-B targets in an overnight 1:10 E:T killing assay (Figure 7D, E). In contrast, all CD8^+^ CAR-T showed similar contact time and cytotoxicity with higher antigen density 691-T tumoroids (Figure 7D-F). Analyzing cytotoxicity by detection of organoid death during the imaging (tumor cell conversion from ‘yellow’ to ‘red’ in Figure 7C) confirmed the observation that TE9 CAR-T were the most rapid at killing B7H3^dim^ 691-B tumoroids followed by 376.96 and MGA271 CAR-T cells. In contrast, all CAR-T cells were equally rapid at killing 691-T tumoroids of higher antigen density (Figure 7G).

There were no differences between binders in terms of the number of new CAR-T contacts during the 800 minute data collection, using either tumoroid model (Supplementary Figure 11A). Hence the finding of highest total T cell/tumoroid contact time of the TE9 CD8 CAR-T cells (Figure 7F) implied TE9 was the binder conferring the longest duration of individual contacts. Consistent with this observation, TE9 CD8^+^ CAR-T cells also showed the lowest displacement distance (Figure 7H) and speed of displacement (Figure 7I). whilst MGA271 had the shortest and 376.96 were intermediate (Figures 7F,H,I). This ranking of tumor interaction duration of TE9 > 376.96 > MGA271 when targeting low antigen density targets correlated with the cytotoxicity, avidity, and binder association rate constant (Supplementary Figure 11B). These data suggest that high avidity binder CD8^+^ CAR-T cells stay longer in contact with low antigen tumoroids to facilitate more efficient killing.

Compared to CD8^+^ CAR-T cells, CD4^+^ CAR-T cells showed greater displacement and reduced tumoroid contact, and this was observed in both models of higher and lower antigen density respectively (Figure 7F,H,I). Moreover, in CD4^+^ CAR-T, the influence of the respective binders on tumoroid contact and displacement was variable, consistent with CD4^+^ CAR-T cells having a lesser role in tumoroid contact and direct cytotoxicity, as described previously using BEHAV3D analysis^24^ (Figure 7F,H,I). While more mobile overall, CD4^+^ CAR-T cells did not engage in CAR-T / CAR-T interaction more than CD8^+^ CAR-T cells (Supplementary Figure 11C).

### Distinct CAR-T cell characteristics revealed by dimensionality reduction of behavioral parameters

To extract behavioral patterns and dynamics, the day 0 BEHAV3D tumor challenge imaging data was clustered using UMAP (Supplementary Figure 12A), revealing six CAR-T behavior types: ‘tumoroid engagers’ (cluster 1), ‘multi-taskers’ (clusters 2, 3), ‘CAR-T engagers’ (clusters 4,5) and ‘dying T cells’ (cluster 6) (Figure 7J). Fitting with the individual cell analysis, the ‘tumoroid engager’ cluster 1 was relatively enriched in CD8^+^ cells compared to CD4^+^ and with higher antigen density 691-T compared with 691-B targets (Figure 7K). Moreover, TE9 and 376.96 binder CD8^+^ T cells consistently exhibited more cluster 1-type tumor engager behavior than MGA271 CAR-T against both tumor targets, and further exhibited more ‘multi-tasker’ behavior in the 691-B tumoroid co-culture. These differences between binders were less apparent in the CD4^+^ CAR-T context.

CAR-T behavior was then compared between the two analyzed timepoints (day 0 and day 7). Day 7 CAR-T behavior was reduced using UMAP into 7 clusters across the two imaging sessions (Supplementary Figure 12B), which grouped into ‘organoid interaction’ (clusters A, B, C, D), ‘CAR-T interaction (clusters E, F) and cell death (cluster G) (Supplementary Figure 12C). Strikingly, no consistent differences between the CAR-T binders or even CD4^+^ *vs* CD8^+^ T cells remained evident at day 7 (Supplementary Figure 12D). This narrowing of CAR-T phenotype was evident when comparing the trajectory of tumoroid death over the day 0 and day 7 imaging sessions, pooling data from all three binders and target types (Figure 7L). A broad range of tumor killing trajectory was seen at day 0 whilst at day 7 all CARs mediated rapid tumor clearance. This apparent ability of low avidity MGA271 CAR-T to control low antigen density tumor at day 7 is most likely due to artificially resetting the effector to target ratio at each tumor challenge with re-sorted viable CAR-T cells.

The dimensionality reduction BEHAV3D analysis reflects the broad conclusion that high avidity of CD8^+^ T CAR-T cells equips them for longer interactions with low antigen density targets, which facilitates sustained cytotoxic activity. The data supports our hypothesis that the differences in CAR-T functionality seen *in vivo* and *in vitro* assays are caused not by variation in binder-driven functional exhaustion, but rather by the differential ability of the CAR-T constructs to proliferate and engage in serial tumor cytotoxicity.

## Discussion

CAR-T cell technology represents a highly promising cancer therapy as exemplified by FDA approvals for several agents in the hemato-oncology field. In solid cancers the pace of development has been slower. The solid cancer environment is immunologically hostile and hypoxic, which limits of capacity of infiltrating CAR-T Cells to engage in the sustained proliferation and cytotoxicity, which is a requirement for shrinking a solid tumor in which the ratio of tumor to CAR-T cells is stacked in favour of the former. CAR-T relative failure is likely compounded by T cell exhaustion, a phenomenon whereby T cells become transcriptionally programmed to hypo-functionality, as a result of repeated antigen stimulation. The capacity to engage in efficient serial killing through repeated successful interactions with tumor cells without inducing T cell exhaustion may be the defining feature of a successful solid cancer CAR-T construct.

The CAR-T to tumor interaction encompasses a CAR immune synapse, wherein the initial contact between T cell and tumor is stabilised by the CAR antibody to target antigen binding interaction, which develops into a cluster of engaged CAR constructs^25,26^. Although the biophysical properties of the CAR-T to antigen synapse are not as well characterized as native TCR immune synapses, it is likely that the strength of this interaction as well as the number of target antigens will determine whether the synapse becomes functional; *i.e.* triggers death of the target cell through release of cytolytic granules concomitant with T cell signalling to promote proliferation and cytokine secretion^25,26^.

Successful formation of a synapse leads to the exclusion of phosphatases (*e.g.* CD45) and the recruitment of src family kinases (*e.g.* lck) resulting in phosphorylation of CD3ζ chain ITAMs within the CAR molecule, and subsequent signalling. An efficient and high-quality CAR-T immune synapse might be one that is strong enough to induce death of the target and short-lived enough to induce a “goldilocks” quantity and quality of T cell signalling that promotes activation without exhaustion. Immune synapse quality is likely affected by the nature of the initiating antibody-to-antigen interaction. Recent progress in CAR-T / target interaction tracking at the single cell level, on platforms such as BEHAV3D^24^, has enabled a nascent understanding of the kinetics of immune synapse formation and how synaptic formation dynamics differ between CD4 and CD8 cells. The relationships between the synapse, the binding properties of the CAR scFv’s and the effect of synaptic signalling on T cell activation and exhaustion remain poorly understood.

Functional avidity is the strength of interaction between a CAR-T cell and its tumor target, which we have measured as the force required to disrupt the interaction. We hypothesized that avidity would be influenced by three factors: the CAR surface expression on CAR-T cells, the target antigen density on the tumor cells, and the affinity of the antibody to antigen interaction at the single molecular interaction scale. We reasoned that this interplay of factors would be such that none of the parameters alone would be predictive of CAR-T response, but the interaction of the three factors to define functional avidity would be the closest predictor of subsequent T cell function. We evaluated the relationship of antibody to antigen biding properties as both affinity (Biacore surface plasmon resonance) and functional avidity (Lumicks) measurements against a range of antigen densities. Two binders of similar affinity (K_D_) had quite differing rate constants of association and dissociation with similar avidity (TE9 and 376.96), whilst a third of lower affinity also demonstrated low avidity (MGA271).

Several previous studies have investigated avidity of CAR-T cells using the Lumicks platform. For example, the group of Maher *et al* studied a range of anti-CD19 binders in the context of leukemia therapy using conventional 2^nd^ generation CARs co-expressing a 4-1BB chimeric costimulatory receptor^27^. Interestingly, intermediate avidity CAR-T were identified to have the greatest *in vivo* capacity to control tumor growth.

Using repeat stimulation assays against suspension cells, monolayers and three dimensional tumoroid models to recapitulate the repeat antigenic engagement of the tumor environment, we show that low avidity interaction on first tumor encounter results in a weak cytokine and proliferative response. Weak initial response translates into ultimate failure of expansion and tumor control in stress conditions. In contrast, higher avidity initial interaction in the low antigen density setting results in effective proliferation and serial killing. In the setting of higher antigen density, all CAR-T constructs have sufficiently high avidity initial interaction to induce expansion and serial killing although low avidity MGA271 still ultimately has reduced functionality at the end of serial challenges. Previous studies have identified fast off-rate as linked to more efficient and effective CAR-T control of leukemia^5,15–17^ and it is therefore an interesting contrast in our tumoroid studies to note that the quickest construct for tumor control of low antigen density had the longest organoid contact time as well as the fastest on– and off-rates. This apparent discordance might be explained by the multiple cell contacts that single CD8^+^ CAR-T generate with the multi-cellular tumoroid structures as previously described using BEHAV3D analysis^24^. Here, a more complex and dynamic interaction of an individual CD8^+^ CAR-T cell with multiple tumor cell targets within a 3D structure might explain a more complex relationship between avidity and success in tumor control.

Of note, in the current models which extend to seven repeat tumor challenges we have not seen evidence of differences in T cell exhaustion between the three binders. This may be because the repeat antigenic challenges were not sustained long enough and it is also tempting to speculate that a higher affinity antibody / higher avidity CAR-T cell might have induced exhaustion in the same repeat stimulation experiments. All binders evaluated showed only modest increases in TIM-3^+^PD-1^+^ double-positive cells and little evidence of exhaustion induced by signalling in the absence of antigen (tonic signalling). It will be important in future studies to evaluate whether very high avidity binders and/or tonic signaling binders show relative failure associated with increased exhaustion marker expression.

One notable observation was that the TE9 binder, which had similar affinity and avidity to the 376.96 binder, had both the highest on-rate and fastest on-rate. To determine if this impacted on the CAR-T / tumor interaction time, we performed BEHAV3D video microscopy analysis. Interestingly, we found that at higher antigen densities there were no difference in dwell time between the binders. In contrast, at low antigen density, the CD8^+^ 376.96 and TE9 CAR-T cells had longer dwell times than MGA271 CAR-T, which was consistent with both their greater cytotoxicity and the higher displacement score of the MGA271 CAR-T. The fast on– and off-rates (in single molecule Biacore affinity studies) of TE9 appears to translate into longer tumoroid contact, and leads us to speculate that avidity measurements may prove to be of greater predictive value for CAR-T synapse formation characteristics. A further surprising result was the short tumour dwell-time of TE9 CD4^+^ cells, which appeared to be of less importance in direct cytotoxicity. Further studies are required to confirm these observations with other binders and targets, as well as mechanistic studies to determine the significance of differences in avidity between CD4^+^ and CD8^+^ CAR-T cells.

B7H3 is an emerging attractive target for cancer immunotherapy due to its bright expression on the surface of a high proportion of cancer cells, its association with poor prognosis, its potential function as an immune regulator, and its relative absence on healthy tissues ^28,29^. Interestingly, its cancer specificity seems to extend beyond cancer cells in the tumor environment since several lines of evidence point to its expression in a variety of tumor infiltrating cells, such as vascular endothelial cells. Hence, the clinical application may extend beyond cancers in which B7H3 has a role as a driver oncogene. Indeed, its vascular endothelial expression might mitigate tumor heterogeneity being a cause of antigen escape variants which has been a significant consideration diminishing enthusiasm for clinical progression with several alternate antigens. All three of the binders evaluated in this study are of clinical relevance and are being evaluated in a range of clinical trials.

There are three important implications of these avidity findings for clinical activity of these respective binders. The first is threshold of antigen density for activation. B7H3 is expressed at low level on some healthy tissues. A lower avidity binder might have optimal activation threshold to distinguish healthy from tumor tissue to avoid on-target off-tumor toxicity. Secondly, the avidity threshold for activation will be affected by the respective signalling characteristics of the CAR. For example, work using the MGA271 binder has shown that substitution of the CD8 hinge and transmembrane regions (H/TM) as evaluated in this paper, for CD28 H/TM, within the context of 41BB-CD3ζ signaling, results is significantly improved function against low antigen density targets^6^. In current clinical trial NCT05474378 (Mackall and colleagues), the configuration is with CD8 hinge/transmembrane with 41BB costimulation. A second recruiting clinical study (SJ-19-0014; St Jude’s, USA; DeRenzo *et al*) using MGA271 with CD28ζ is deploying an additional 41BBL co-stimulation *in-trans* module^22^, whilst trial NCT04185038^14^ (Seattle, USA; Vitanza and team) makes use of MGA271 with CD28 H/TM in a 2^nd^ gen 41BB-CD3ζ configuration. Hence, none of the current MGA271 trials are using the configuration in the current paper. Thirdly, higher expression of CAR through use of higher MOI of virus could increase avidity of a given binder against targets. Artificially, we ensured equivalent expression levels to allow fair comparison of avidity effects in relation to target antigen density. Higher expression of CAR might increase avidity and functionality. What seems clear is that all CARs require fine-tuning of binders to identify optimal activation thresholds. Avidity measurements may prove an additional valuable parameter for screening constructs to optimal signalling and activation.

## Acknowledgements

We thank Debarati Shome and Reza Nadafi of Lumicks for their support with our z-Movi analysis of B7H3-CAR-T synapse avidities, and Lumicks more broadly for supporting us with hardware and technical advice during the experiments. We’re grateful to the Flow Cytometry Core Facility at UCL GOS Institute of Child Health and to the Imaging Center at the Princess Máxima Center for their support in running our studies. We thank the Princess Máxima Center Imaging and Tumoroid facilities for support.

## Funding

We are grateful for the following grant support. Anderson and Chesler: Stand up to Cancer/ CRUK Paediatric Cancer New Discoveries Challenge; Anderson, Chesler and Donovan: CwC INSTINCT-MB award, Fight Kids Cancer, and LPT programme awards; Anderson: Research into Childhood Cancer, NIHR GOSH BRC, GOSHCC 552864, AMR/LifeArc awards. Drost and Buhl: Oncode Institute, Children Cancer-free Foundation (KiKa), Deutsche Forschungsgemeinschaft (German Research Foundation, DFG). Molenaar: Villa Joep, Veni. This work has received funding from the European Union’s Horizon 2020 research and innovation program under the Marie Skłodowska–Curie grant agreement, No. 956285 (VAGABOND).

## Author Contributions

MB, JW, and JA wrote the manuscript; MB, HM, EZ, MBR, and SbD analyzed data; GF generated reagents; EZ, HM, RS, BD, CH, CB, SMT, MN, SM, KB, CLB, AV, LP, KoS, AG, and MB performed experiments; JA, JW, JB, KC, AR, JD, JM, LC co-supervised the research.

## Conflicts of Interest

MB, JA, KC, KB, MB hold patents in CAR-T technology development, including a pending patent for the TE9 anti-B7H3 binder. JA holds founders shares in Autolus.

## Methods

### Cells lines and culture conditions

All cell lines were cultured in humidified 37C° in 5% CO_2_ conditions. Cell lines were thawed and passaged for at least two weeks prior to use in functional assays. Culture mycoplasma tests were performed monthly using MycoAlert Detection Kit from Lonza. The following table summarizes the characteristics and culture conditions of the cell lines that were used over the course of this study.

| *Cell Line ID* | *Type of Cancer* | *Source* | *RRID* | *Culture Media* |
| --- | --- | --- | --- | --- |
| *LAN-1 (LA-N-1)* | Neuroblastoma | ECACC | CVCL_1827 | RPMI1640 (Gibco) + 10% FCS (Gibco) |
| *Med8A (MED-MEB-8A) wild type* | Medulloblastoma | Gift from Michael Taylor, The Hospital for Sick Children, Canada | CVCL_M137 | DMEM (Gibco) + 10% FCS + 1x Glutamax (Gibco) |
| *Med8A GFP/Luc* | Medulloblastoma | Derived from Med8A wild type by RD114 γ-retroviral transduction with GFP/Luc | None | DMEM (Gibco) + 10% FCS + 1x Glutamax (Gibco) |
| *Kelly* | Neuroblastoma | Gift from Andrew Stoker, UCL, UK | CVCL_2092 | RPMI1640 + 10% FCS |
| *SupT1 wild type* | Childhood T lymphoblastic lymphoma | ECACC | CVCL_1714 | RPMI1640 + 10% FCS |
| *SupT1-B7H3<sup>hi</sup>* | Childhood T lymphoblastic lymphoma | Derived from SupT1 wild type by RD114 γ-retroviral transduction with 4-Ig B7H3 (Sinobiological), followed by flow cytometric sorting | None | RPMI1640 + 10% FCS |
| *SupT1-B7H3<sup>range</sup> GFP/Luc* | Childhood T lymphoblastic lymphoma | Derived from SupT1 wild type by RD114 γ-retroviral transduction with GFP/Luc followed by flow cytometric sorting, then 4-Ig B7H3 (Sinobiological) without sorting | None | RPMI1640 + 10% FCS |
| 691-T<br>(AMC691-T)<br>wild type | Neuroblastoma | Gift from Jan Molenaar<br>(Utrecht) | None | 'Neurosphere medium': DMEM-GlutaMAX supplemented with 20% Ham's F-12 nutrient mixture, 2% B-27 supplement minus vitamin A, 1% N-2 Supplement, 100 U/mL penicillin, 100 mg/mL streptomycin, 300 ng/mL insulin-like growth factor (IGF-1), 40 ng/mL fibroblast growth factor-basic (FGF-2), 20 ng/mL epidermal growth factor (EGF), 10 ng/mL platelet-derived growth factor-AA (PDGF-AA) and 10 ng/mL platelet-derived growth factor-BB (PDGF-BB) |
| 691-T<br>(AMC691-T)<br>GFP/Luc | Neuroblastoma | Gift from Jan Molenaar<br>(Utrecht) | None | 'Neurosphere medium' |
| 691-B<br>(AMC691-B) | Neuroblastoma | Gift from Jan Molenaar<br>(Utrecht) | None | 'Neurosphere medium' |
| 691-B<br>(AMC691-B)<br>GFP/Luc | Neuroblastoma | Gift from Jan Molenaar<br>(Utrecht) | None | 'Neurosphere medium' |
| 103-T wild<br>type | Malignant<br>rhabdoid tumor | Gift from Jarno Drost<br>(Utrecht) | None | 'MRT medium'<sup>30,31</sup>: MRT tumoroids were cultured in droplets of growth factor-reduced basement membrane extract (BME) Type 2 (R&D Systems) in kidney tumoroid medium (KOM) (Advanced Dulbecco's Modified Eagle Medium (DMEM)/F12 (Gibco) containing 1X GlutaMAX (Thermo Fisher Scientific), 10 mM HEPES (Thermo Fisher Scientific) and 1X Penicillin-Streptomycin (P/S) (Merck Millipore), supplemented with 10% R-spondin-conditioned medium, 1.5% B27 supplement (Gibco), 50 ng/mL epidermal growth factor (EGF) (PeproTech), 50 ng/mL fibroblast growth factor (FGF)-2 (PeproTech), 1.25 mM N-acetylcysteine (Sigma), 10 μM Rho-associated coiled-coil containing protein kinase (ROCK) inhibitor Y-27632 (Abmole) and 5 μM A83-01 (Tocris Bioscience)) |
| 103-T Luc | Malignant<br>rhabdoid tumor | Gift from Jarno Drost<br>(Utrecht); derived from 103-T wild type by lentiviral transduction with pLKO.1-UbC-luciferase blast and pLV-H2B-mNeon-puro cassettes<sup>32,33</sup> | None | 'MRT medium' |
| D341 wild<br>type | Medulloblastoma | Gift from Michael Taylor,<br>The Hospital for Sick<br>Children, Canada | CVCL_0018 | DMEM (Gibco) + 10% FCS + 1x Glutamax (Gibco) |
| *D341 GFP/Luc* | Medulloblastoma | Derived from D341 wild type by RD114 γ-retroviral transduction with GFP/Luc | None | DMEM (Gibco) + 10% FCS + 1x Glutamax (Gibco) |
| *D425 wild type* | Medulloblastoma | Gift from Michael Taylor, The Hospital for Sick Children, Canada | CVCL_1275 | DMEM (Gibco) + 10% FCS + 1x Glutamax (Gibco) |
| *D425 GFP/Luc* | Medulloblastoma | Derived from D425 wild type by RD114 γ-retroviral transduction with GFP/Luc | None | DMEM (Gibco) + 10% FCS + 1x Glutamax (Gibco) |
| *293-T (HEK293T)* | Transformed cell line | ATCC | CVCL_0063 | IMDM + 10% FCS |

### CAR sequences

All CARs were expressed using an SFG backbone in a 2^nd^ generation RD114 γ-retroviral system. All plasmids were sequenced to verify absence of unexpected sequence mutations prior to manufacture of virus. A diagram of construct structure is shown in Figure 1A, with all anti-B7H3 CARs expressed in the following format: RQR8-T2A-scFv-CD8αLinker-CD8αTM-CD28endodomain-CD3ζchain. The sequences for the scFv’s were obtained from public sources and are summarized in the below table. To insert these into the CAR backbone, the scFv sequences were ordered as geneblocks and then restriction cloned into a CD8α-CD28-CD3ζ format.

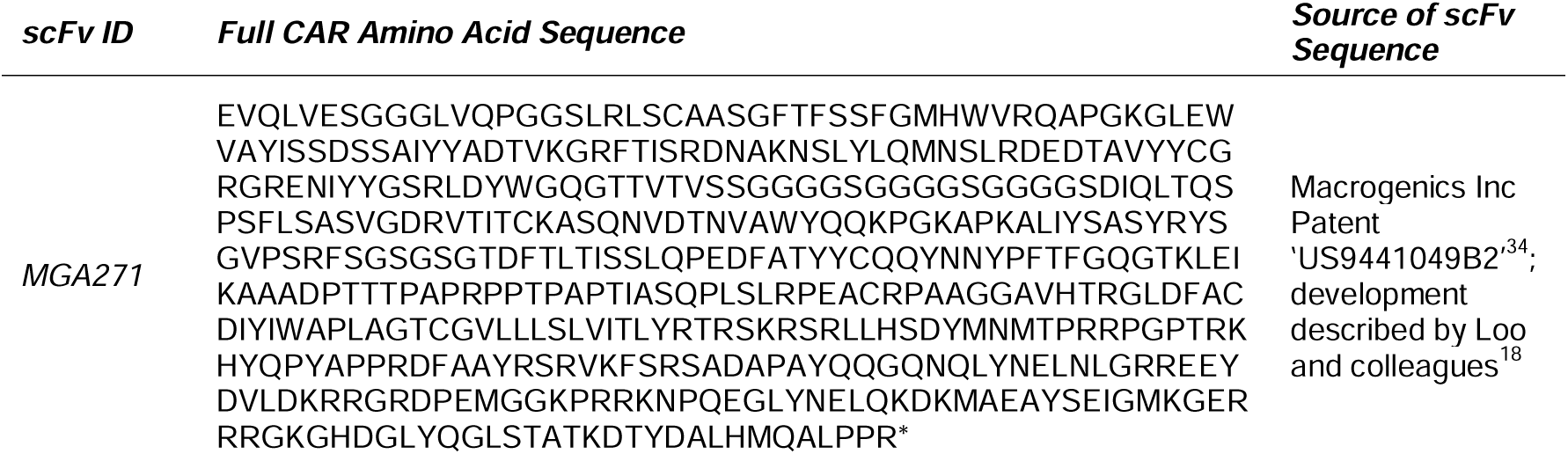
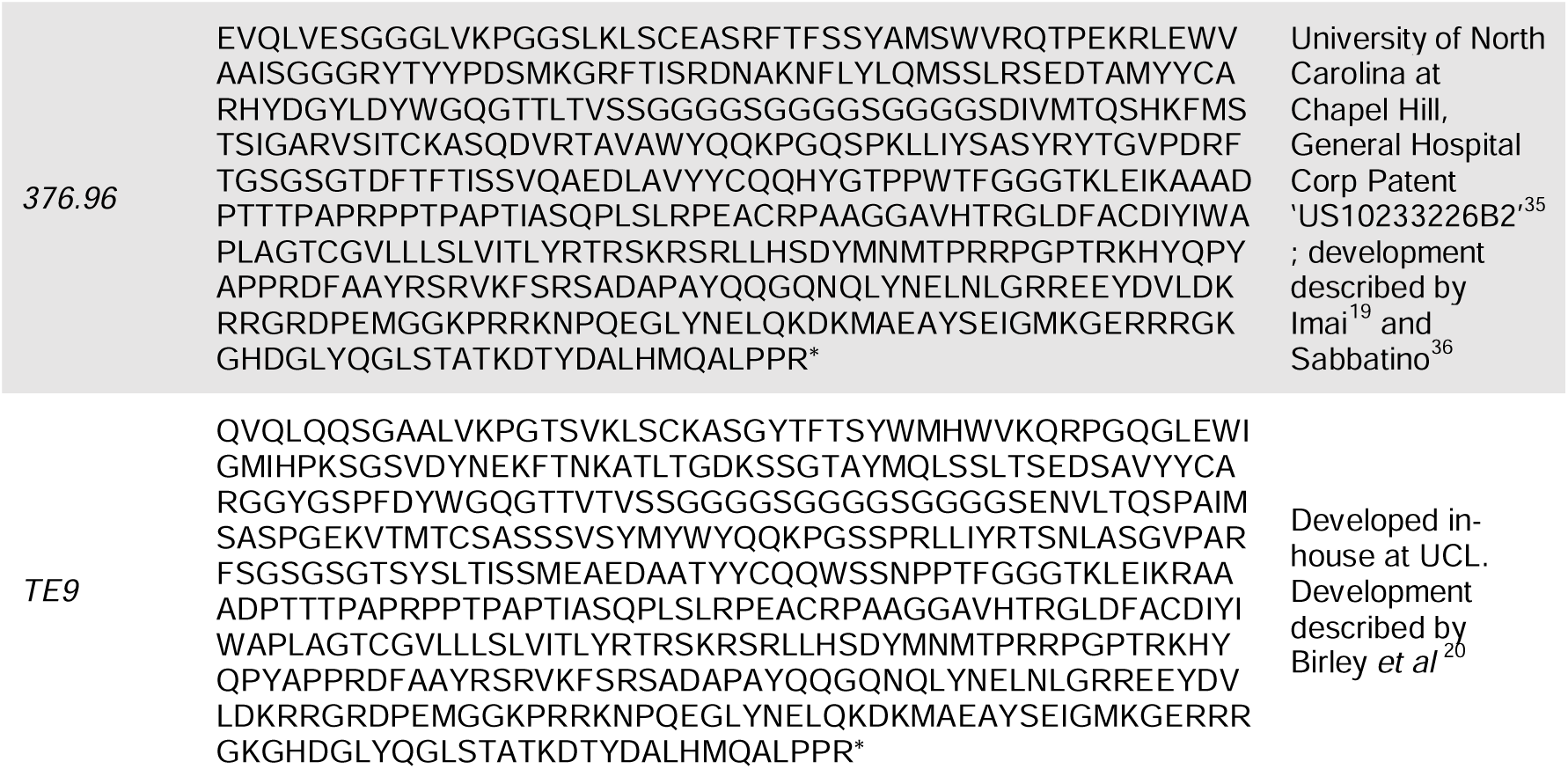

### CAR-T cell manufacture

Leukapheresis cones were acquired from NHS Blood and Transplant. PBMCs were separated through Ficoll centrifugation using Lymphoprep (STEMCELL Technologies). PBMCs were washed and residual red cells lysed with ACK Lysis buffer (Thermo Fisher Scientific). NK cells were depleted using magnetic CD56 depletion beads (Miltenyi Biotec) and LD depletion columns (Miltenyi Biotec). PBMCs were suspended in RPMI1640 containing 10% FCS and 1x L-glutamine at a concentration of 1×10^6^ cells/mL, and then activated with 0.5 mg/mL of anti-CD3 (clone: OKT-3; Miltenyi Biotec) and anti-CD28 antibodies (clone: CD28.2; Miltenyi Biotec). Forty-eight hours before transduction and on the day of transduction, 100 IU/mL recombinant human IL-2 (Proleukin, Novartis) was added. T cells were transduced with the above-mentioned γ-retroviral SFG construct following plating on retronectin (Takara Bio)-coated plates and spinoculation. Transduction efficiency was measured three days after transduction using flow cytometry by staining for RQR8 (clone: QBNED10; R&D BioSystems). T cell populations were not corrected for transduction efficiency in functional assays. Transduction efficiencies of >50% were routinely achieved. Expanding CAR-T cell IL-2 media was replenished every two days. Expansions were harvested at day 10-12 following culture initiation and cryopreserved in 10% DMSO complete media. When initiating *in vitro* or *in vivo* functional assays, CAR-T cells were thawed in pre-warmed IL-2 (100 IU/mL) complete media (RPMI1640, 10% FCS), rested overnight and then counted. CAR-T E:T ratios were based on the number of viable T cells after overnight rest. Cryopreserved CAR-T recovery after overnight rest (relative to the number of CAR-T that were frozen down) was routinely 50-60%, with a ∼90% viability.

### Animal studies

Animal protocols were approved by local institutional research committees and in accordance with UK Home Office guidelines. Mixed male and female NSG mice aged between 6 and 8 weeks were supplied by Charles River. All experiments were carried out under UK home office licenses project license number 15981/01, and personal license numbers I13398879, P2E645DD9, I83870811, I27052867, I8C20AD0D and I754094BC.

For the LAN-1 *in vivo* study, NSG mice were injected with 1 x 10^6^ LAN-1-BFP/Luc in Matrigel (Corning) subcutaneously into the flank. CAR-T cells (1 x 10^7^) were injected intravenously into the tail vain at day 10 following tumor engraftment (treatment schedule illustrated in Figure 1E). Tumor size was monitored twice a week with digital callipers. Mice were given 200 uL luciferin (Thermo Fisher Scientific) into the scruff and imaged using a PhotonIMAGERTM optical imaging system (Biospace Lab) weekly. Animal sacrifice was initiated upon reaching either a tumor diameter of >15 mm in any direction (as measured by callipers) or the appearance of tumor burden toxicity that exceeds our animal welfare standards (>15% weight loss, change in activity, ataxia, dehydration, loss of normal gait, grimacing).

For the Med8A *in vivo* study, two types of model were run: the ‘high-stress’ model and the ‘low-stress’ model. In both models 5 x 10^4^ Med8A cells re-suspended in PBS were injected into the cerebellum of an NSG mouse. In the ‘low-stress model’ (illustrated in Figure 1B), two days after tumor engraftment the mice were administered intraventricularly with 2.5 x 10^6^ CAR-T cells. In the ‘high-stress model’ (illustrated in Figure 1C) the mice were treated six days after tumor engraftment, and with 0.5 x 10^6^ CAR-T cells per animal. Mice were given 200 uL luciferin (Thermo Fisher Scientific) into the scruff and imaged using a PhotonIMAGERTM optical imaging system (Biospace Lab) weekly. Animal sacrifice was initiated upon reaching either a tumor luminescence of 10E10 photons/second/cm^2^ (via IVIS), or the appearance of tumor burden toxicity that exceeds our animal welfare standards (>15% weight loss, doming of the cranium, change in activity, ataxia, dehydration, loss of normal gait, grimacing).

### CAR-T in vitro functional assays

A number of different functional CAR-T assays were performed. While high (>60%) transduction efficiency of CAR-T was routinely achieved, transduced cell numbers were normalised for functional assays to ensure that each binder condition receives the same number of viable CAR-T cells. All data was collected in experimental duplicates or triplicates, as indicated in figure legends.

CAR-T responses against LAN-1 and Kelly target cells were assessed as follows. For the overnight (∼18h) co-culture cytokine assays, CAR-T cells were co-cultured with LAN-1, Kelly cells or no antigen stimulus in 48-well plates at an E:T ratio of 1:1. After 18 h, supernatant was removed for ELISA and cells incubated with monensin (BioLegend). Checkpoint receptors PD-1, TIM-3 and LAG-3 were detected by flow cytometry. Overnight cytotoxicity was tested using a ^51^Cr release assay. Target cells were incubated with ^51^Cr for 1 h then washed and plated in 96-well plates. CAR-T cells or untransduced cells were plated at a range of E:T ratios. The plates were incubated for 4 h at 37°C and then the supernatant was removed and incubated with scintillation fluid (PerkinElmer) overnight at room temperature. ^51^Cr released into the supernatant was measured using a 1450 MicroBeta TriLux (PerkinElmer). For the re-challenge assays, CAR-T cells were co-cultured with irradiated LAN-1, Kelly, or no target cells in 24-well plates at an E:T ratio of 2:1. Cell medium was replenished every 2–3 days. CAR-T cells were challenged with irradiated target cells every 6 days, cultured for a further 24 h, and analyzed. Cells were pelleted and supernatant was removed every week for ELISA. CAR-T cell proliferation was measured weekly by flow cytometry using Precision Count Beads (BioLegend). The levels of cytokines IL-2 and IFN-γ were quantified using ELISA MAX Deluxe Set Human IL-2 and ELISA MAX Deluxe Set Human IFN-γ (BioLegend).

SupT1-B7H3^range^ re-challenge assays were performed as follows (illustrated in Figure 6C). For the overnight (∼18h) co-culture cytokine assays, CAR-T cells were co-cultured with GFP/Luc-SupT1-B7H3^range^ cells or GFP/Luc-SupT1-WT cells in 96-well plates at an E:T ratio of 1:1. After 18 h, supernatant was removed for analysis with ELISA MAX Deluxe Set Human IL-2 and ELISA MAX Deluxe Set Human IFN-γ (BioLegend). Cytotoxicity at the various indicated timepoints was tested using a luciferase-based assay. Briefly, 15ug/mL of D-luciferin (Perkin Elmer) was added to each well to document the remaining number of living tumor cells. The plate was then incubated for 5 min at 37°C in 5% CO_2_ and run on a SpectraMax, Molecular Devices analyzer, with ‘no effector’ tumor wells as controls for T cell-mediated tumor death. CAR-T cells or untransduced cells were plated at a 1:1 E:T ratio, with daily re-challenges. Cytotoxicity readings were done in this manner 18h after each new tumor re-stimulation. CAR-T cell proliferation was measured weekly by flow cytometry using Precision Count Beads (BioLegend).

Brain tumor challenge assays with GFP/Luc-Med8A, –D425 and –D341 cells were done by plating CAR-T cells with targets at the indicated E:T ratios in 96-well plates. Cytotoxicity at 96h following tumor challenge was tested using a luciferase-based assay. Briefly, 15ug/mL of D-luciferin (Perkin Elmer) was added to each well to document the remaining number of living tumor cells. The plate was then incubated for 5 min at 37°C in 5% CO_2_ and run on a SpectraMax, Molecular Devices analyzer, with ‘no effector’ tumor wells as controls for T cell-mediated tumor death. CAR-T cell proliferation was measured weekly by flow cytometry using Precision Count Beads (BioLegend).

Malignant rhabdoid tumoroids were established, characterised and cultured as previously described^30,31^. 103-T tumoroids were transduced with a construct encoding for luciferase were cultured in KOM in BME droplets for 7 days in advance. 4 days before the co-culture the tumoroids were disrupted mechanically into a single cell suspension by mechanical dissociation and seeded in BME droplets. On the day of CAR-T challenge, MRT tumoroids were collected, washed with cold medium to remove BME, and a fraction was dissociated into a single cell suspension. Single tumor cells were counted to determine the number of cells present in the MRT tumoroid suspension. MRT tumoroids were then seeded out with an equivalent of 7500 single tumor cells per well in 50 uL of co-culture medium containing 10% FBS and a 1:1 ratio of KOM and RPMI with 1X GlutaMAX and 1X P/S. To determine tumor specific killing, a released luciferase assay was performed using the luciferase assay system (Promega). Medium was removed and cells were washed with 100 uL of PBS. 20 uL of 1X passive lysis buffer (Promega) were added and the plate was incubated at room temperature for 15 min while shaking. In a LUMITRAC white polystyrene 96-well plate (Greiner) 100 uL of the luciferase reagent were added to 20 uL of the lysed cells and the plate was measured at the FluoSTAR Omega Microplate reader immediately. Luciferase signal was determined on day 1 after co-culture as well as before re-challenge with new tumoroids.

To measure CAR-T cytotoxicity against (AMC) 691-B and 691-T neuroblastoma tumoroids, GFP-Luc tumoroids were dissociated into single cells by re-suspending in 1 mL of Accutase solution (Sigma-Aldrich) through a p10 tip on top of a p1000 tip to combine both enzymatic and mechanic dissociation methods. Tumoroid cells in suspension were counted and plated two days before the beginning of the experiment to ensure spheroid formation in 96-well white flat bottom plates(Costar). On the day of initiation co-culture, CAR-T cells were counted and plated with GFP/Luc tumoroids at the stated E:T ratios in half ‘neurosphere medium’ (expanded on in the Cell Lines section of Methods and Materials) and half T cell medium (RPMI-GlutaMAX 1640 + 25 mM Hepes (ThermoFisher) supplemented with 10% FCS (Gibco), 1% penicillin/streptomycin (ThermoFisher) and 1% L-glutamine (ThermoFisher) supplemented with 100 IU/mL IL-2 (Miltenyi Biotec)). For re-challenge, tumoroids were plated 2 days before each challenge, with the CAR-T culture added on top. Tumoroid re-challenge a set E:T ratio was done every 2-3 days. Cytotoxicity was measured by adding 15ug/mL of D-luciferin (Perkin Elmer) to each well to document the remaining number of living tumor cells, with ‘no effector’ tumor wells as controls for T cell-mediated tumor death. Cells were then incubated for 5’ at 37°C in 5% CO_2_ and readout performed in the FLUOstar Omega microplate reader.

### Flow cytometry staining and analysis

The following cell cytometers were used to collect flow cytometry data: BD ® LSR II Flow Cytometer (BD Biosciences), FACSymphony™ A5 High-Parameter Cell Analyzer (BD Biosciences) and CytoFLEX S V4-B2-Y4-R3 Flow Cytometer (Beckman Coulter). FlowJo 10.9.0 software for Mac (Becton Dickinson) was used to analyse flow cytometry data. All staining was done in FACS buffer, PBS, 2% FCS with 2mM EDTA. Samples were either analysed immediately following staining or fixed with Biolegend Fixation Buffer and analysed within two weeks. The following table lists the flow cytometry staining reagents that were used:

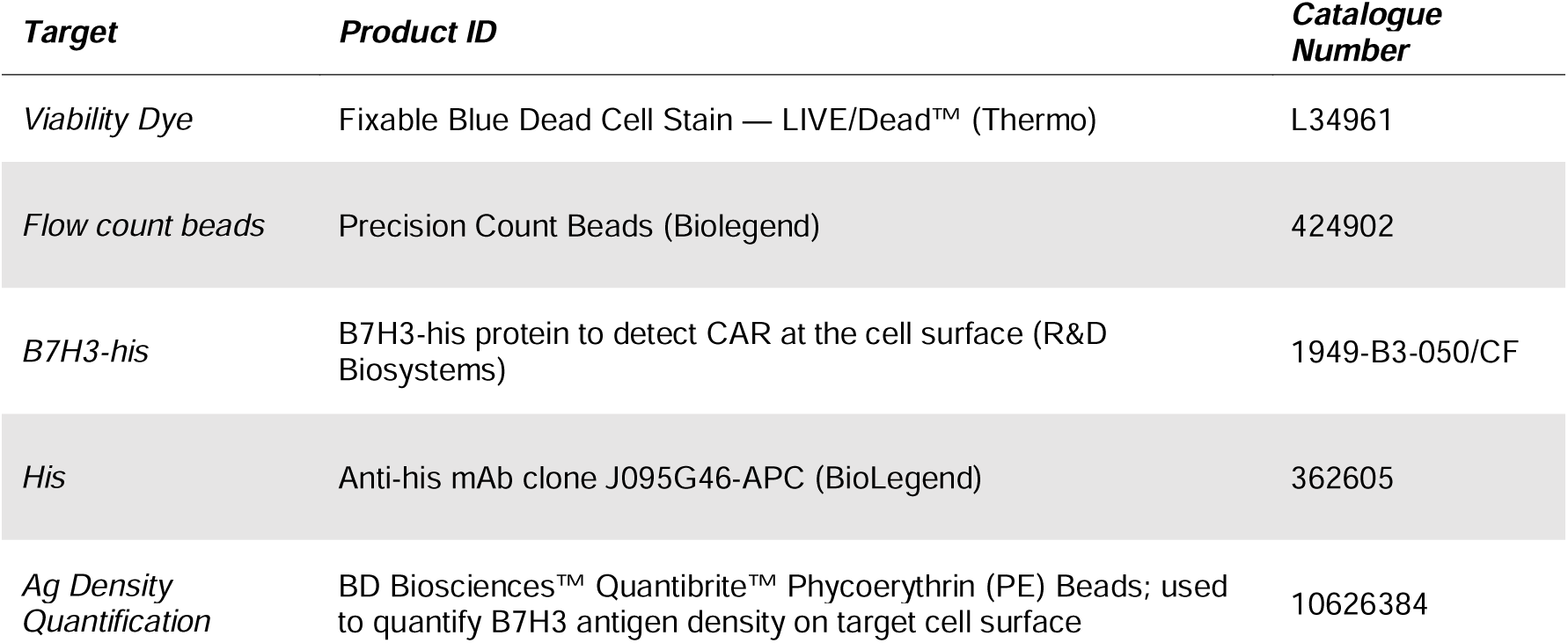
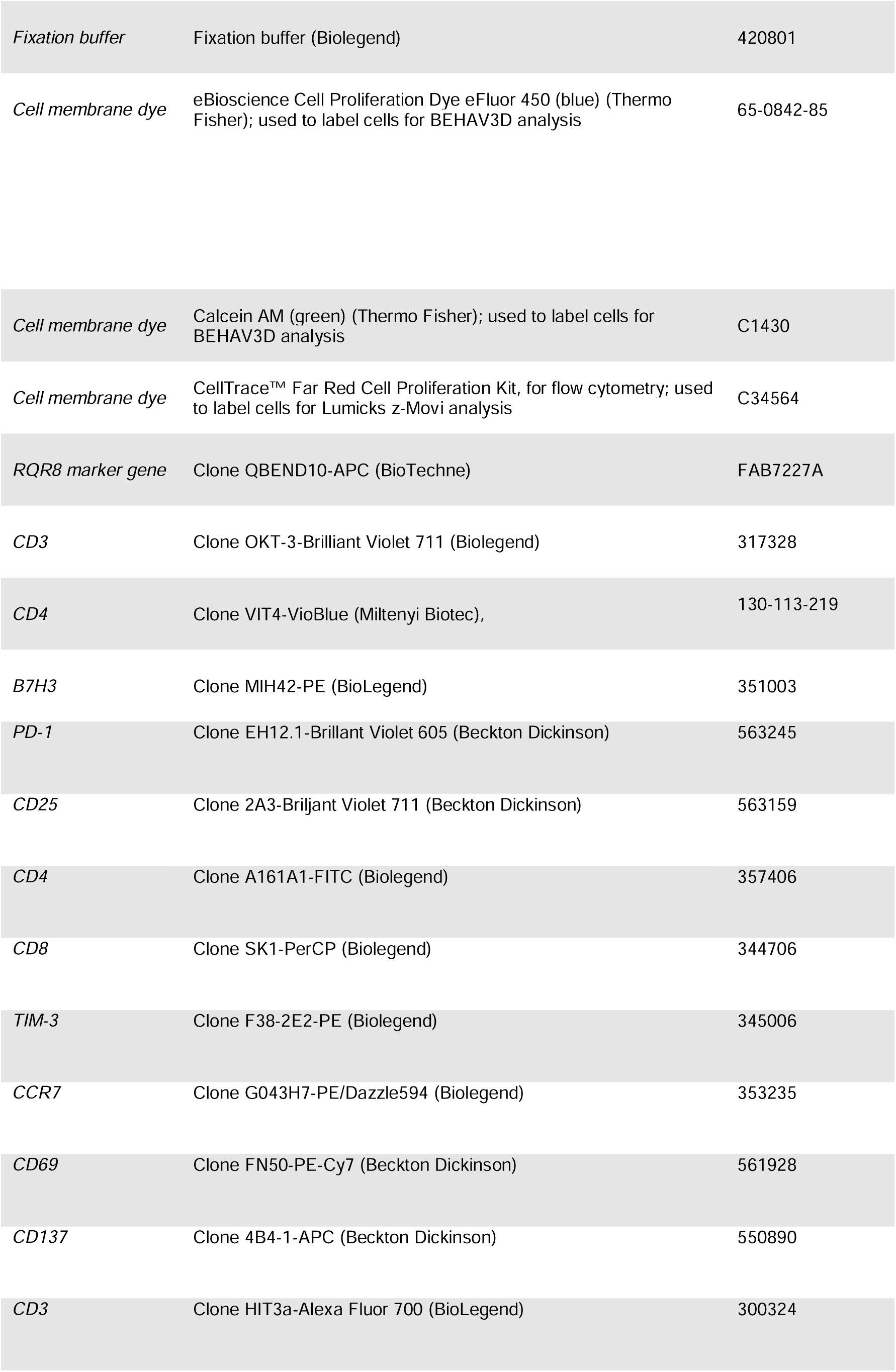
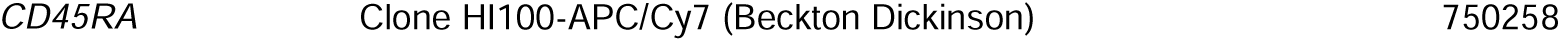

### Lumicks CAR-T synapse avidity measurements

Proprietary z-Movi Lumicks protocols were followed under the supervision of Lumicks staff present for the assays. Briefly, SupT1-WT, SupT1-B7H3^hi^, LAN-1 and Med8A tumor cells were seeded in a z-Movi microfluidic chip (Lumicks, Amsterdam, Netherlands) coated with poly-L-lysine and cultured for 16 hours. The next day, flow sorted, transduction efficiency-normalized and CellTrace Red (Biolegend)-labelled CAR-T cells were serially flowed in the chips and incubated with the target cells for 10 minutes prior to initializing a 3-minute linear force ramp. During the force ramp, the z-Movi device (Lumicks) captured a time series of images using a bright field microscope integrated into the platform. Detached cells were levitated towards the acoustic nodes, allowing the tracking of cells based on their XY positions. Changes in the Z-position resulted in a change in the diffraction pattern, which allowed the distinction between cells adhered to the substrate and cells suspended to the acoustic nodes. This information was used to correlate cell detachment events with a specific rupture force. Cell detachment was acquired using z-Movi Tracking and post experiment image analysis was done using Cell Tracking offline analysis. Data were presented as ‘mean % CAR-T attached cells’ at the maximum mean force applied, 1000 piconewtons, pN. This was repeated in experimental triplicate for four biological T cell donors. A greater depth of z-Movi workflow is reported online^37^. The software used to acquire the data was z-Movi Software (v1.0).

### Binder affinity measurements

Biacore surface plasmon resonance (SPR) for anti-B7H3 antibodies was done as previously described^38^. Briefly, MGA271, 376.96 and TE9 binders were expressed in human IgG1 format (Evitria), and then evaluated by Antibody Analytics using Biacore SPR for binding kinetics to 4-Ig human B7H3. Anti-B7H3 antibodies were first adhered to plates and several concentrations of Human B7-H3 analyte were passed over the surface to assess the binding interactions. The data is shown in Figure 2G. Clone 9G8 mAb was used as an isotype control.

### BEHAV3D CAR-T / tumoroid imaging

Tumoroids (691-B and –T) were grown in ‘Neurosphere medium’ (as described in the ‘Cell Lines’ section), cultured for 3 weeks prior to the experiments and passaged once or twice a week according to tumoroid size and confluence. The experimental timeline is illustrated in Figure 7A. CAR T cells were thawed one day prior to the start of the experiment in RPMI-GlutaMAX 1640 + 25 mM Hepes (ThermoFisher) supplemented with 10% FCS (Gibco), 1% penicillin/streptomycin (ThermoFisher) and 1% L-glutamine (ThermoFisher), rested overnight at a density of 1×10^7^ cells/mL in 100 IU/mL IL-2 (Miltenyi Biotec)-supplemented medium. CAR T cells for the initial challenge day 0 imaging experiment were stained with the following antibody mix in FACS buffer (PBS 1x + 2% FCS + 2mM EDTA): anti-RQR8 mAb clone QBEND10-APC (R&D Systems), anti-CD4 mAb clone VIT4-VioBlue (Miltenyi Biotec) and mouse serum (Invitrogen). RQR8^+^CD4^+^ and RQR8^+^CD4^-^ CAR-T cells were FACS-sorted with a Sony SH800S cell sorter into FCS, and rested overnight in a density of 1×10^7^ cells/mL in medium supplemented with 100 IU/mL IL-2 (Miltenyi Biotec).

For co-culture experiments, tumoroids were dissociated into single cells by pipetting 1 mL of Accutase solution (Sigma-Aldrich, counted and plated two days before the beginning of the experiment to ensure spheroid formation in either 12-well flat-bottom plates (for rec-hallenges) (Greiner) or in a glass-bottom 96-well SensoPlate (Greiner) for imaging. The correct cell number was determined considering organoid growth rates. Wild type organoids were used. Sorted and rested CAR T cells were stained separately with eBioscience Cell Proliferation Dye eFluor 450 (1:4000, ThermoFisher) or Calcein AM (1:3000, ThermoFisher) in PBS for 15 minutes at 37°C and mixed in a 1:1 ratio immediately before plating them with the organoids in a 1:5 E:T ratio. The co-culture medium (100μL organoids + 100μL T cells) was then supplemented with 2.5% basement membrane extract (BME, Cultrex), Nucred Dead 647 (three drops per mL, ThermoFisher), and TO-PRO-3 (1:5000, ThermoFisher). The combination of Nucred Dead 647 and TO-PRO-3 has previously been described by Dekkers et al^24^. The plate was placed in a SP8 confocal microscope (Leica) containing an incubation chamber (37°C, 5% CO2) and imaged for thirteen hours in time series with two-minute intervals (as shown in Supplementary Figure 10). Day 7 BEHAV3D imaging was the third challenge with 691 tumoroids. On Day 6, cells from re-challenge plates were stained with the following antibody mix in FACS buffer (PBS1x + 2% FCS + 2mM EDTA): anti-RQR8 mAb clone QBEND10-APC (R&D Systems), anti-CD4 mAb clone VIT4-VioBlue (Miltenyi Biotec), anti B7H3 mAb clone MIH42-PE (BioLegend) and mouse serum (Invitrogen). B7H3^-^ CD34^+^CD4^+^ and B7H3^-^ CD34^+^CD4^-^ CAR-T cells were FACS-sorted with the Sony SH800S cell sorter into FCS, rested overnight at a density of 1×10^7^ cells/mL in medium supplemented with 100 IU/mL IL-2 (Miltenyi Biotec). On day 7, rested CAR-T cells were stained separately and plated with WT organoids in a 1:5 E:T ratio, as described above. The plate was placed in a SP8 confocal microscope (Leica) containing an incubation chamber (37°C, 5% CO2) and imaged for thirteen hours in time series with a two-minute interval.

Image processing was done according to the protocol previously described by Dekkers *et al*^24^ using the Imaris (Oxford Instruments) version 10.0 software for 3D visualization, cell segmentation and extraction of statistics. The Channel Arithmetics Xtension was used to create additional channels for the specific identification of CD8^+^ and CD4^+^ T cells (live and dead), organoids (live and dead), and to exclude debris. The Surface and ImarisTrack modules were used to specifically detect and track T cells and/or organoids. For tracked T cells, time-lapse statistics consisting of the coordinates of each cell, speed, square displacement, distance to either organoids or the other T cell subset, and dead cell dye channel intensity were exported into a metadata file for subsequent processing and analysis.

### Data presentation

Data was graphed and analysed statistically using Graphpad Prism 10 for macOS software (Dotmatics). Statistical tests were chosen as suitable to particular data sets, and statistical test recommendations by Graphpad Prism 10 software were followed, based on data type, replicates and degree of pairing. The number of biological and experimental replicates used in each analysis is listed in respective figure legends. Data processing for the BEHAV3D platform was done using ‘R’ programming language code that was written by and published from the Rios group^24^.

## Notes

### Summary of Updates

additional funding information added to the end of the acknowledgement section; SPecific funding for Vagabond PhD programme has been added

## References

1. Research, C. for B. E. and. Approved Cellular and Gene Therapy Products. FDA (2023).

2. Albelda, S. M. CAR T cell therapy for patients with solid tumours: key lessons to learn and unlearn. Nat Rev Clin Oncol 21, 47–66 (2024).

3. Weber, E. W. et al. Transient “rest” restores functionality in exhausted CAR-T cells via epigenetic remodeling. Science 372, eaba1786 (2021).

4. Weber, E. W. et al. Pharmacologic control of CAR-T cell function using dasatinib. Blood Adv 3, 711–717 (2019).

5. Ghorashian, S. et al. Enhanced CAR T cell expansion and prolonged persistence in pediatric patients with ALL treated with a low-affinity CD19 CAR. Nature Medicine 25, 1408–1414 (2019).

6. Majzner, R. G. et al. Tuning the Antigen Density Requirement for CAR T-cell Activity. Cancer Discov 10, 702–723 (2020).

7. Du, H. et al. Antitumor Responses in the Absence of Toxicity in Solid Tumors by Targeting B7-H3 via Chimeric Antigen Receptor T Cells. Cancer Cell 35, 221–237.e8 (2019).

8. Theruvath, J. et al. Locoregionally administered B7-H3-targeted CAR T cells for treatment of atypical teratoid/rhabdoid tumors. Nat Med 26, 712–719 (2020).

9. Majzner, R. G. et al. CAR T Cells Targeting B7-H3, a Pan-Cancer Antigen, Demonstrate Potent Preclinical Activity Against Pediatric Solid Tumors and Brain Tumors. Clinical Cancer Research 25, 2560–2574 (2019).

10. Maachani, U. B. et al. B7-H3 as a Prognostic Biomarker and Therapeutic Target in Pediatric central nervous system Tumors. Transl Oncol 13, 365–371 (2020).

11. Modak, S., Kramer, K., Gultekin, S. H., Guo, H. F. & Cheung, N. K. Monoclonal antibody 8H9 targets a novel cell surface antigen expressed by a wide spectrum of human solid tumors. Cancer Res 61, 4048–4054 (2001).

12. Chapoval, A. I. et al. B7-H3: A costimulatory molecule for T cell activation and IFN-γ production. Nature Immunology 2, 269–274 (2001).

13. Wang, L., Kang, F.-B. & Shan, B.-E. B7-H3-mediated tumor immunology: Friend or foe? International Journal of Cancer 134, 2764–2771 (2014).

14. Vitanza, N. A. et al. Intraventricular B7-H3 CAR T Cells for Diffuse Intrinsic Pontine Glioma: Preliminary First-in-Human Bioactivity and Safety. Cancer Discov 13, 114–131 (2023).

15. Chmielewski, M., Hombach, A., Heuser, C., Adams, G. P. & Abken, H. T Cell Activation by Antibody-Like Immunoreceptors: Increase in Affinity of the Single-Chain Fragment Domain above Threshold Does Not Increase T Cell Activation against Antigen-Positive Target Cells but Decreases Selectivity1. The Journal of Immunology 173, 7647– 7653 (2004).

16. Mause, E. R. V. et al. Combinatorial T cell engineering eliminates on-target off-tumor toxicity of CD229 CAR T cells while maintaining anti-tumor activity. 2021.12.06.471279 Preprint at 10.1101/2021.12.06.471279 (2022).

17. Olson, M. L. et al. Low-affinity CAR T cells exhibit reduced trogocytosis, preventing rapid antigen loss, and increasing CAR T cell expansion. Leukemia 36, 1943–1946 (2022).

18. Loo, D. et al. Development of an Fc-enhanced anti-B7-H3 monoclonal antibody with potent antitumor activity. Clin Cancer Res 18, 3834–3845 (2012).

19. Imai, K., Wilson, B. S., Bigotti, A., Natali, P. G. & Ferrone, S. A 94,000-dalton glycoprotein expressed by human melanoma and carcinoma cells. J Natl Cancer Inst 68, 761–769 (1982).

20. Birley, K. et al. A novel anti-B7-H3 chimeric antigen receptor from a single-chain antibody library for immunotherapy of solid cancers. Molecular Therapy – Oncolytics 26, 429–443 (2022).

21. Philip, B. et al. A highly compact epitope-based marker/suicide gene for easier and safer T-cell therapy. Blood 124, 1277–1287 (2014).

22. Nguyen, P. et al. Route of 41BB/41BBL Costimulation Determines Effector Function of B7-H3-CAR.CD28ζ T Cells. Mol Ther Oncolytics 18, 202–214 (2020).

23. Labanieh, L. et al. Enhanced safety and efficacy of protease-regulated CAR-T cell receptors. Cell 185, 1745–1763.e22 (2022).

24. Dekkers, J. F. et al. Uncovering the mode of action of engineered T cells in patient cancer organoids. Nat Biotechnol 1–10 (2022) doi:10.1038/s41587-022-01397-w.

25. Sajman, J. et al. Nanoscale CAR Organization at the Immune Synapse Correlates with CAR-T Effector Functions. Cells 12, 2261 (2023).

26. Xiong, Y., Libby, K. A. & Su, X. The physical landscape of CAR-T synapse. Biophysical Journal (2023) doi:10.1016/j.bpj.2023.09.004.

27. Halim, L. et al. Engineering of an Avidity-Optimized CD19-Specific Parallel Chimeric Antigen Receptor That Delivers Dual CD28 and 4-1BB Co-Stimulation. Frontiers in Immunology 13, (2022).

28. Kontos, F. et al. B7-H3: An Attractive Target for Antibody-based Immunotherapy. Clinical Cancer Research 27, 1227–1235 (2021).

29. Zhou, W.-T. & Jin, W.-L. B7-H3/CD276: An Emerging Cancer Immunotherapy. Frontiers in Immunology 12, (2021).

30. Schutgens, F. et al. Tubuloids derived from human adult kidney and urine for personalized disease modeling. Nat Biotechnol 37, 303–313 (2019).

31. Calandrini, C. et al. An organoid biobank for childhood kidney cancers that captures disease and tissue heterogeneity. Nat Commun 11, 1310 (2020).

32. Custers, L. et al. Somatic mutations and single-cell transcriptomes reveal the root of malignant rhabdoid tumours. Nat Commun 12, 1407 (2021).

33. Koo, B.-K. et al. Controlled gene expression in primary Lgr5 organoid cultures. Nat Methods 9, 81–83 (2011).

34. Johnson, L. S., Huang, L., Moore, P. A., Loo, D. T. & Chen, F. Z. Antibodies reactive with B7-H3 and uses thereof. (2014).

35. Dotti, G., Ferrone, S., Du, H., Wang, X. & Ferrone, C. Methods and compositions for chimeric antigen receptor targeting cancer cells. (2019).

36. Sabbatino, F. et al. Novel Tumor Antigen-Specific Monoclonal Antibody-Based Immunotherapy to Eradicate Both Differentiated Cancer Cells and Cancer-Initiating Cells in Solid Tumors. Seminars in Oncology 41, 685–699 (2014).

37. 37. The z-Movi Cell Avidity Analyzer workflow. *LUMICKS* https://lumicks.com/knowledge/the-z-movi-workflow/.

38. Jason-Moller, L., Murphy, M. & Bruno, J. Overview of Biacore systems and their applications. Curr Protoc Protein Sci Chapter 19, Unit 19.13 (2006).

